# Functional integration and segregation during semantic cognition: Evidence across age groups

**DOI:** 10.1101/2023.11.02.565291

**Authors:** Wei Wu, Paul Hoffman

## Abstract

Semantic cognition is underpinned by ventral anterior temporal lobe (vATL) which encodes knowledge representations and inferior frontal gyri (IFG), which controls activation of knowledge based on the needs of the current context. This core semantic network has been validated in substantial empirical findings in the past. However, it remains unclear how these core semantic areas dynamically communicate with each other to achieve successful semantic processing. Here, we investigated this question by testing functional connectivity in the core semantic network during semantic tasks and whether these connections were affected by cognitive ageing. Compared to a non-semantic executive task, semantic tasks increased the connectivity between left and right IFGs, indicating a bilateral semantic control system. Strengthened connectivity was also found between left IFG and left vATL, and this effect was stronger in the young group. At a whole-brain scale, IFG and vATL increased their coupling with multiple-demand regions during semantic tasks, even though these areas were deactivated relative to non-semantic tasks. This suggests that the domain-general executive network contributes to semantic processing. In contrast, IFG and vATL decreased their interaction with default mode network (DMN) areas during semantic tasks, even though these areas were positively activated by the task. This suggests that DMN areas do not contribute to all semantic tasks: their activation may sometimes reflect automatic retrieval of task-irrelevant memories and associations. Taken together, our study characterized a dynamic connectivity mechanism supporting semantic cognition within and beyond core semantic regions.

## Introduction

Semantic cognition refers to our ability to acquire conceptual knowledge about the world and use this knowledge to generate task and context-appropriate behaviours (Hoffman, McClelland, & Lambon Ralph, 2018; Jefferies, 2013; Lambon Ralph, Jefferies, Patterson, & Rogers, 2017). Successful semantic cognition is critical to normal function in daily activities across the lifespan, due to its central role in bringing meaning to verbal and non-verbal stimuli. The controlled semantic cognition (CSC) framework (Jefferies, 2013; Lambon Ralph et al., 2017) proposes that semantic cognition is underpinned by two distinct but dynamically interactive elements: semantic knowledge representations, which encode the meanings and properties of concepts (e.g., words and objects), and semantic control processes, which are responsible for the retrieval and manipulation of this knowledge to guide our behaviour.

Converging evidence indicates that semantic knowledge representations are encoded in a distributed set of brain regions that overlaps partially with the default mode network (Binder & Desai, 2011; Binder, Desai, Graves, & Conant, 2009; Devereux, Clarke, Marouchos, & Tyler, 2013; Fernandino et al., 2016; Jackson, 2021), while semantic control processes engage a left-lateralised frontotemporal network that overlaps partially with domain-general executive systems (i.e., the multiple demand network; Jackson, 2021; Noonan, Jefferies, Visser, & Lambon Ralph, 2013; Quillen, Yen, & Wilson, 2021; Rodd, Davis, & Johnsrude, 2005; Vitello & Rodd, 2015; Whitney, Kirk, O’Sullivan, Lambon Ralph, & Jefferies, 2011). Within these networks, two specific regions, which we focus on in the present study, appear to play critical, specific roles in semantic cognition. The inferior frontal gyrus (IFG) plays a central role in controlled semantic processing (Badre & Wagner, 2007; Hoffman, Jefferies, & Lambon Ralph, 2010; Thompson-Schill, D’Esposito, Aguirre, & Farah, 1997; Vitello & Rodd, 2015), while the ventral part of anterior temporal lobe (vATL) forms the centrepoint of a multimodal knowledge representation hub (Binney, Embleton, Jefferies, Parker, & Ralph, 2010; Mion et al., 2010; Rice, Hoffman, & Lambon Ralph, 2015; Rice, Lambon Ralph, & Hoffman, 2015; Rogers et al., 2021). While the functions of these regions have been separately investigated in detail, another crucial theoretical claim of the CSC has received less attention. The CSC proposes that semantic knowledge and control regions dynamically interact with each other to generate semantically-appropriate behaviour. Despite their importance, the nature of these interactions is less studied. Examining these interactions are also central to understanding semantic cognition in later life, when semantic knowledge is well-developed but controlled processing declines (Hoffman, 2018; Spreng & Turner, 2019; Turner & Spreng, 2015). Accordingly, the current study used functional connectivity analyses across different semantic tasks to investigate three questions:

1. How do functional connections between vATL and IFG, and between these regions and broader neural networks, change when people engage in semantic processing?
2. How are functional connections between left IFG/vATL and their right-hemisphere homologues influenced by semantic processing?
3. Where do these effects differ between young and older people?

IFG is a critical region in semantic processing, but the specific contributions of left and right IFGs have been a long-standing mystery. Left IFG plays a central role in semantic control processes: convergent evidence shows that the left IFG is activated during semantic tasks and reliably shows increased activation as semantic demands increase (Hoffman et al., 2010; Jackson, 2021; Jung, Rice, & Lambon Ralph, 2021; Krieger-Redwood, Teige, Davey, Hymers, & Jefferies, 2015; Lambon Ralph et al., 2017; Quillen et al., 2021). However, the functional role of the right IFG in semantic cognition is unclear. The right IFG sometimes shows activation during semantic tasks but to a lesser degree than the left IFG (Jackson, 2021; Krieger-Redwood et al., 2015). Its activation seems to be enhanced in particular populations. Right IFG exhibits increased activation in aphasic patients following left-hemisphere stroke, but debate continues as to whether this activity is beneficial for language processing or is maladaptive (Gainotti, 2015; Stefaniak, Halai, & Lambon Ralph, 2020; Turkeltaub, 2015). Healthy older people also activate the right IFG more than young participants during semantic tasks, particularly when their performance is impaired (for meta-analysis, see Hoffman & Morcom, 2018). Thus, the major debate over the right IFG’s activation in semantic tasks is whether this region supports the processing of the dominant left IFG (e.g., when semantic demands are high or when the left IFG’s function is compromised due to damage or age); or whether right IFG activation instead reflects a failure to suppress functionally-irrelevant cortex, a phenomenon frequently referred to as “neural dedifferentiation” in the cognitive aging literature (Li, Brehmer, Shing, Werkle-Bergner, & Lindenberger, 2006; Li & Rieckmann, 2014; Morcom & Henson, 2018; Park & Reuter-Lorenz, 2009; Reuter-Lorenz & Cappell, 2008; Reuter-Lorenz & Park, 2014; Spreng & Turner, 2019).

Uncertainty about the functional role of right IFG has led to this area being neglected in models of semantic processing. To rectify this, it is necessary to test how this region interacts with the left IFG and other core semantic regions. In a recent study, we examined the left and right IFGs’ activation profiles along parametrically-varied semantic demands in healthy young and older people (Wu & Hoffman, 2023). Both IFGs activated during semantic tasks, though the left IFG showed stronger activation than the right. Importantly, there was a consistent linear demand- activation relationship in both IFGs across age groups and semantic tasks: both IFGs showed activation increases as task difficulty increased (though the effect was stronger in the left). These results suggest that right IFG plays a functional role in semantic processing, supporting the dominant left IFG. The current study builds on these findings by investigating functional connectivity of right IFG. If the right IFG assists the left IFG in controlled semantic processing, we would expect to see increased coupling between these two areas during semantic tasks. We would also expect both IFGs to show similar patterns of increased connectivity with other semantic regions.

The functional roles of left and right ATLs are also a matter of ongoing debate. Unlike IFG, there is widespread agreement that both ATLs contribute to semantic processing. However, the degree of functional specialisation between them is disputed. While some models propose segregated representations in left and right ATLs for verbal and non-verbal knowledge (Gainotti, 2012, 2014; Snowden, Thompson, & Neary, 2004), others suggest a more integrated view in which the two ATLs jointly contribute to representation (Lambon Ralph, Cipolotti, Manes, & Patterson, 2010; Rice, Lambon Ralph, et al., 2015). While arguing for a generally bilateral representation system, the latter view proposes that left ATL comes to exhibit relative specialisation for language semantics, by virtue of its stronger connections with left-lateralised speech production and orthographic systems (Hoffman & Lambon Ralph, 2018; Schapiro, McClelland, Welbourne, Rogers, & Lambon Ralph, 2013).

How the left and right ATLs interact with each other during semantic tasks has rarely been investigated. In one study that combined fMRI with transcranial magnetic stimulation (TMS), researchers found that the activation of right vATL increased when the left vATL was inhibited by stimulation, and that the right vATL increase was positively correlated with semantic task performance (Jung & Lambon Ralph, 2016). In another study, researchers found the functional connectivity between left and right vATLs increased when semantic tasks became more demanding (Jung et al., 2021). Finally, patients with surgical resection of the left ATL show more activation in right vATL than controls during semantic tasks (Rice, Caswell, Moore, Lambon Ralph, & Hoffman, 2018). These findings are compatible with a bilateral, co-operative model of ATL semantic representation, in which right vATL contributions increase when demands increase or left vATL function is compromised. However, the evidence so far is limited and is mostly based on data from natural or virtual lesions.

Beyond left-right connectivity in the same regions, there is also a strong theoretical drive to understand how the knowledge-related ATL and semantic-control IFG interact with each other. The CSC framework claims that this cross-region interaction is the means by which we use our knowledge to generate appropriate behaviours in varied contexts (Hoffman et al., 2018; Jefferies, 2013; Lambon Ralph et al., 2017). How has the interaction between ATL and IFG been investigated? One commonly used method is to focus on one of the core semantic regions and use it as the seed area to conduct whole-brain level connectivity analysis, revealing the interaction between the seed and other regions. For example, Jackson, Hoffman, Pobric, and Lambon Ralph (2016) investigated how left vATL connects with the other regions in the brain during resting state and during an active semantic task. They found overlapped connectivity patterns across the two contexts, in which the left vATL showed significant connectivity with a range of semantic-related areas such as IFG and medial prefrontal cortex. For IFG, Chiou, Humphreys, Jung, and Lambon Ralph (2018) investigated how the interaction between the left IFG and knowledge-related regions changes in different semantic tasks. Compared to a typical semantic association task (e.g., pairing *ketchup* with *mustard*), a task which required selective extraction of colour knowledge (e.g., pairing *ketchup* with *fire extinguishers*) enhanced the connectivity between the left IFG and occipitotemporal cortex, a visual knowledge representation area. In another study, researchers have compared the whole-brain functional connectivity patterns elicited by a semantic task versus a visual perception task (Chiou, Jefferies, Duncan, Humphreys, & Lambon Ralph, 2023). They found the seed area in left IFG showed more connectivity with the executive multiple demand network (MDN) regions during the semantic task and more connectivity with the default mode network (DMN) during the visual task.

The above studies have made significant contributions to understanding how the core semantic areas connect with other regions in the brain at a broad topographic level. However, they did not quantify how the core semantic regions (i.e., vATL and IFG) change their coupling with each other during semantic processing. In a limited number of cases where the ROI-to-ROI connectivity in the semantic network has been examined (for example, see Chiou et al., 2018; Jung et al., 2021), analyses have mostly targeted vATL and IFG connectivity in the left hemisphere without considering cross-hemisphere interactions or the roles of right IFG or right vATL. In addition, studies commonly use tasks that place high demands on controlled semantic processing (e.g., attending selectively to particular semantic features). These tasks presumably require strong interaction between semantic control and knowledge areas. In contrast, tasks which place demands on knowledge representations themselves (e.g., understanding low-frequency vocabulary) may engage the semantic system in a different way.

To address the above questions, the current study systematically investigated functional connectivity between left and right IFGs and vATLs in two different semantic tasks, which either required high levels of semantic control or placed high demands on knowledge representations. As well as testing connections between these regions, we also used each region as a seed to explore its whole-brain connectivity pattern during semantic tasks. Finally, we tested how the coupling of semantic regions is influenced by healthy ageing. Recently, a domain-general theory of cognitive ageing has been proposed named ‘the default-executive coupling hypothesis of aging’ (DECHA) (Spreng & Turner, 2019; Turner & Spreng, 2015). DECHA proposes that, compared to young people, older adults rely more on their crystallized intelligence (e.g., knowledge obtained from prior experience) and less on their fluid intelligence (e.g., executive control) when completing cognitive tasks. DECHA suggests that this shift in cognitive architecture is accompanied by stronger but less flexible interaction between knowledge representation areas and executive control regions.

The DECHA account predicts that older people are less able to decouple knowledge and control areas when prior knowledge is not relevant to the current task (e.g., in a visual perception task). The consequences for semantic cognition are less clear, as here interaction between knowledge and control systems is likely to be beneficial for performance. In a recent study, we examined these effects in the domain of semantic cognition at a broad network level (Wu & Hoffman, 2023). Contrary to DECHA’s predictions, we did not find that older people showed less flexible connectivity between knowledge-related DMN and either the executive control MDN or the semantic control network (SCN). However, although the vATL and IFG are embedded in the above networks, these networks also include a range of other regions that do not necessarily contribute to semantic processing. It is possible that the DECHA connectivity predictions are more applicable to specific core semantic regions rather than more distributed neural networks. We used the current study to test this possibility and to explore how semantic-related connectivity differs between age groups.

## Materials and Methods

The current study uses fMRI data collected by Wu and Hoffman (2023). The dataset is publicly available (https://doi.org/10.7488/ds/3845 and https://doi.org/10.7488/ds/3846).

### Participants

fMRI scans were acquired for 45 older adults and 45 young adults, who were recruited from the Psychology department’s volunteer panel and local advertising, and participated in the study for payment. All participants were native English speakers, reported to be in good health with no history of neurological or psychiatric illness. The older participants completed the Mini-Addenbrooke’s Cognitive Examination (M-ACE; Hsieh et al., 2015) as a general cognitive screen. Two older participants were excluded because they scored < 26 of 30 on the M-ACE. One older participant was excluded because we were unable to identify seed co-ordinates in the right vATL for them. Two young participants’ data were also excluded due to technical issues or structural abnormalities. In the end, data from 42 older participants (27 females, 15 males; mean age = 67.93 years, s.d. = 5.09 years, range = 60 - 79) and 43 young participants (31 females, 12 males; mean age = 23.07 years, s.d. = 3.23 years, range = 18 - 32) were used in the analyses. Both age groups had a high level of formal education (older adults: mean = 15.62 years, s.d. = 2.87 years, range = 10 - 22; young adults: mean = 17.07 years, s.d. = 2.53 years, range = 12 - 23), and young adults had completed more years of education than older adults (*t* _83_ = 2.47, two-tailed *p* < 0.05). This was expected given that younger generations have greater access to higher education. Informed consent was obtained from all participants. This research was performed in accordance with all relevant guidelines/regulations and the study was approved by the University of Edinburgh Psychology Research Ethics Committee.

## Materials

There were three tasks in the current study: a semantic control test, a semantic knowledge test and a cognitively demanding non-semantic test (see Figure 1 for examples). All task stimuli were obtained from the norms of Wu and Hoffman (2022).

**Figure 1.**
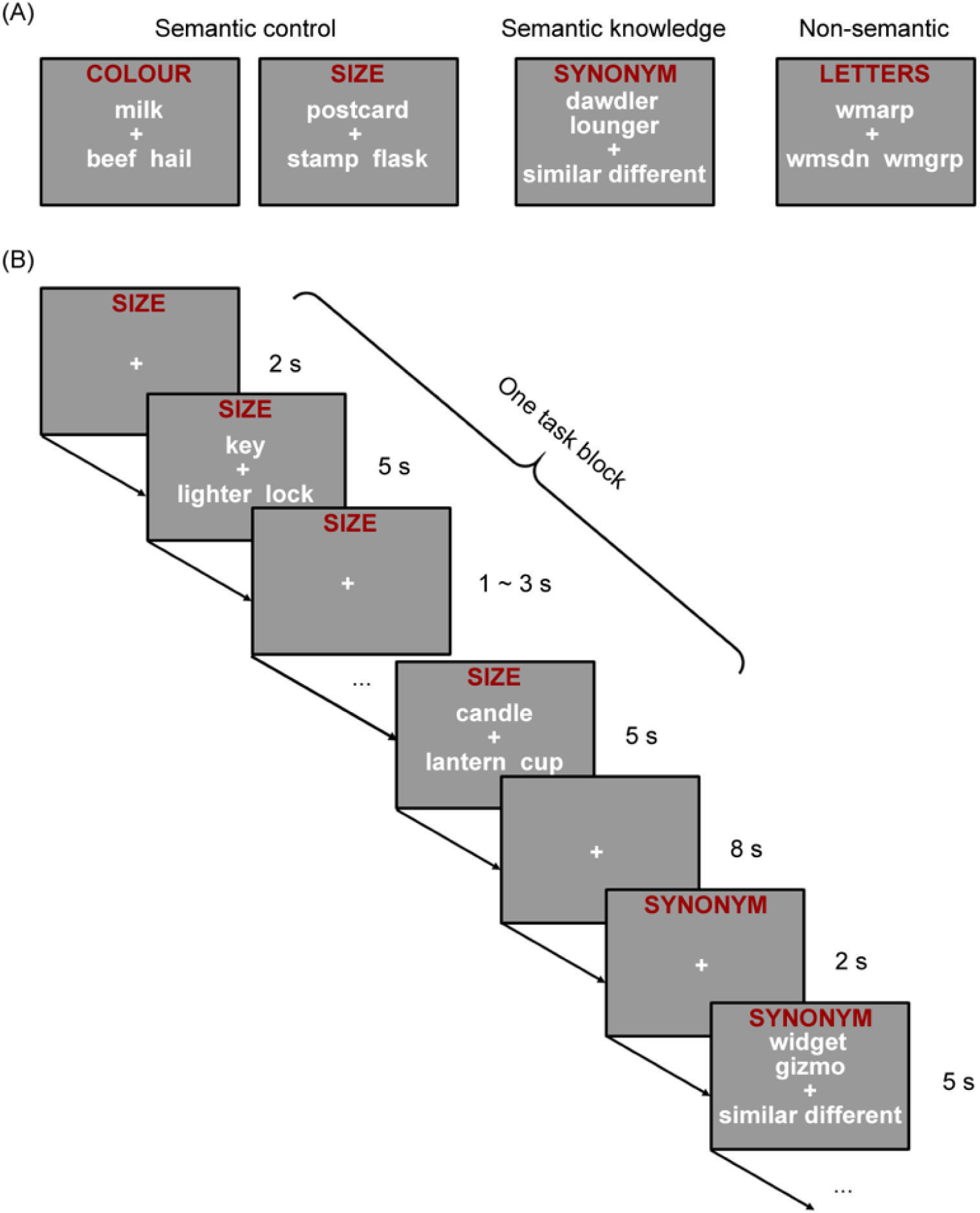
(A) Example items from each task and (B) an illustration of block structure in the experiment.

### Test of semantic control

An 80-trial feature-matching task was completed to probe semantic control processes. On each trial, a probe word above the centre of the screen was shown, along with two option words in a line below. Participants were asked to decide which option shared the same particular semantic feature with the probe (colour on 40 trials and size on 40 trials). For instance, on a colour trial, *milk* would match with *hail* as both are typically white. This task taxes semantic control processes as participants need to direct their attention to the target semantic properties and inhibit competing but irrelevant semantic knowledge and associations from the distractor (e.g., *milk - beef*) (Badre, Poldrack, Pare-Blagoev, Insler, & Wagner, 2005; Thompson-Schill et al., 1997).

### Test of semantic knowledge

Participants completed an 80-trial synonym judgement task designed to tax their semantic knowledge. On each trial, participants were presented with a pair of words above the centre of the screen vertically with the two options (i.e., similar or different) in a line below. They were asked to decide if the two words shared a similar meaning or not. As this task includes lower-frequency words whose meanings are less familiar to people, this task placed high demands on semantic knowledge representations (Wu & Hoffman, 2022).

### Test of non-semantic cognitive control

We used an 80-trial non-semantic task as the baseline task in comparison with the semantic tasks. In this task, each trial was composed of three meaningless letter strings, which were presented in a similar fashion to the feature-matching task. Participants were asked to choose the option that shared the most letters in the same order as the probe. This task required participants to resolve competition between options based on orthographic similarity but without engaging semantic processing. Thus it examined general cognitive control processes.

#### Design and procedure

There were two scanning runs for each participant. In each run, participants were presented with 10 task blocks from each of the three tasks (i.e., 30 blocks in total). In each block, a task cue (i.e., “synonym”, “colour”, “size”, or “letters”) was first shown for 2 s and remained at the top of the screen during the whole block. Then participants viewed four 5-s task trials from the same task, with inter-trial interval lasting for 1-3 s (jittered, 2 s on average). Stimuli in each task were grouped into 5 levels of task difficulty and each block contained four trials from a single difficulty level. The analyses reported here combined across all levels of difficulty. The blocks were separated by 8-s periods of fixation and the order of the blocks in each run was randomized. We also pseudo-randomized trial order within each block so that the position of correct responses in the block (i.e., left or right) was balanced. To counterbalance the potential influences of block/trial order on our data, we produced two sets of experimental programs with different block/trial orders and used each set for half of the participants in each age group. During the experiment, participants used their left and right index fingers to press buttons to indicate their choice. Each task trial was shown once during the whole experiment. For the feature-matching task, all the trials in a block required participants to make judgements based on the same feature (colour or size).

#### Image acquisition and processing

Images were acquired on a 3T Siemens Skyra scanner with a 32-channel head coil. We employed a whole-brain multi-echo acquisition protocol to minimize the impact of head movements and signal drop out in the ventral temporal regions (Kundu et al., 2017). For the functional images, the multi-echo EPI sequence included 46 slices covering the whole brain with echo time (TE) at 13 ms, 31 ms and 50 ms, repetition time (TR) = 1.7 s, flip angle = 73°, 80 × 80 matrix, reconstructed in-plane resolution = 3 mm × 3 mm, slice thickness = 3.0 mm and multiband factor = 2. Two runs of 642 volumes were acquired in total. A high-resolution T1-weighted structural image was also acquired for each participant using an MP-RAGE sequence with 1 mm isotropic voxels, TR = 2.5 s, TE = 4.4 ms. Images were pre-processed and analysed using SPM12 and the TE-Dependent Analysis Toolbox (Tedana) (DuPre et al., 2021). The first 4 volumes of each run were discarded. Estimates of head motion were obtained using the first BOLD echo series. Slice-timing correction was carried out and images were then realigned using the previously obtained motion estimates. We used Tedana to combine the three echo series into a single time series and to divide the data into either BOLD-signal or noise-related. The noise components of the data were discarded. Then images were unwarped with a B0 fieldmap to correct for irregularities in the scanner’s magnetic field. Finally, functional images were spatially normalised to MNI space using SPM’s DARTEL tool (Ashburner, 2007) and were smoothed with a kernel of 8 mm FWHM.

After pre-processing, we first built a general linear model (GLM) to analyse the data from the two experimental runs for each participant, with the data being treated with a high-pass filter with a cut-off of 180 s. In the GLM, there were three regressors for the three different tasks (i.e., semantic control task, semantic knowledge task and non-semantic task). Each trial was modelled as a 5-s event and convolved with the canonical hemodynamic response function. The GLM also included 12 nuisance regressors modelling movement artifacts, which were measured using the three translations and three rotations estimated during spatial realignment, and their scan-to-scan differences. We computed one-sample t-tests over first-level whole-brain contrast maps across all participants, which contrasted each of the semantic tasks versus the non-semantic task, to obtain semantic control and knowledge activation effect maps at a group level. Two-sample t-tests over the individual-level contrast maps were then computed to examine the activation differences between older and young people during semantic processing. We also contrasted model estimates of the average of the two semantic tasks vs the non-semantic task to define seed areas for the PPI analyses.

#### Definition of regions of interest (ROIs)

To define the seed areas in bilateral vATLs and IFGs (Figure 3), we used a method that combined anatomical masks with selection of functionally-activated peak coordinates at the individual participant level. First, anatomical vATL and IFG masks were obtained. The IFGs were defined using the BA 45 mask in the Brodmann Areas Map in MRIcron (https://people.cas.sc.edu/rorden/mricro/template.html). The anatomical vATLs were defined in a similar fashion to a previous study (Hoffman & Lambon Ralph, 2018). Specifically, we generated the fusiform gyrus mask using the voxels with a greater than 50% probability of falling within fusiform gyrus in the LONI Probabilistic Brain Atlas (LPBA40), and we divided the fusiform gyrus mask into 5 roughly-equal-length sections that ran along an anterior-to-posterior axis. The anatomical masks of vATLs were then constructed by combining the first two sections (anterior parts) of the above mask.

Second, at the group level, we used the contrast of the average of the semantic knowledge and control tasks minus the non-semantic task to identify voxels that were most responsive to semantic processing (relative to domain-general cognitive demands). We overlapped this functional map with the anatomical masks to obtain group-level peak coordinates in the vATLs and IFGs (shown in Figure 3). Finally, to identify the peak activation locations for each individual participant, four 15-mm-radius spheres were built around the group-level peaks. We searched within these spheres to identify the peak coordinates for each participant (using the individual-level contrast maps of the average of semantic tasks versus non-semantic task). Individual-level seeds with 8-mm radius for each participant were built centred on these coordinates.

#### Psychophysiological interaction (PPI) analyses

Using PPI, we examined how semantic processing (relative to non-semantic processing) modulated the connectivity between the left and right vATL and IFG areas, as well as between each of these ROIs and the rest of the brain. We performed generalised psychophysiological interactions (gPPI) to achieve this goal (McLaren, Ries, Xu, & Johnson, 2012). PPI analysis tests how the correlation of the activity between two brain areas (i.e., the physiological effect) changes when a participant engages in different contexts or tasks (i.e., the psychophysiological interaction). While traditional PPI can only search for task-specific changes in connectivity between two conditions, gPPI has the advantage that it can include more than two task conditions in a single PPI model, as is required by the current investigation. We used the gPPI toolbox (https://www.nitrc.org/projects/gppi) to perform the PPI analyses.

For each participant and each seed area, we built a gPPI model with the same GLM structure as the earlier contrast analysis, which modelled the three tasks. Additionally, the time courses from the individual-level seed mask was also extracted and added to the gPPI model, along with its interaction with the task regressors (i.e., the PPI terms). Once estimated, this model gave us whole-brain effect maps corresponding to each regressor, in which the maps for PPI terms estimated how the connectivity between the seed region and each voxel in the brain changed for each task relative to fixation. By contrasting the model estimates of the PPI terms, we were able to reveal how connectivity varied with task (e.g., semantic knowledge PPI minus non-semantic PPI). We conducted our further analyses based on these effect maps.

### ROI-level analysis

In this analysis, we were interested in the functional interactions between the core semantic areas in both hemispheres (i.e., left and right vATLs and IFGs) during semantic processing. Thus, for each participant, we used the effect maps obtained from the PPI model (with one ROI as the seed) to extract the effects of each PPI term from the other individual-level ROI masks. For example, a model with left IFG as the seed was used to generate whole-brain PPI effect maps for the contrast of semantic knowledge minus non-semantic tasks. Using this effect map, we extracted the voxel-averaged effects from each of the other ROIs (i.e., right IFG, left vATL and right vATL), which indicated how the correlations between the left IFG and the other ROIs were affected by the semantic knowledge task. As PPI effects are not directional (O’Reilly, Woolrich, Behrens, Smith, & Johansen-Berg, 2012), we used each region in each pair of ROIs once as seed and once as target, and averaged the effects to give a measure of PPI connectivity. Taking the left IFG – left vATL connectivity as an example, this means that we had one model that used the timecourses of the left IFG as the seed to extract effects from the left vATL mask (the target), and we had another model that used the timecourses of the left vATL as the seed to extract effects in left IFG. The averaged estimates from the two models were used to represent the PPI connectivity between left IFG and left vATL. After signal extraction for the contrasts of each semantic task versus the non-semantic task, one-sample t-tests were computed over the PPI effects across all participants and semantic tasks to determine if connectivity varied between general semantic processing versus non-semantic processing. A 2 (older and young groups) by 2 (semantic control and semantic knowledge tasks) ANOVA was further conducted to test connectivity differences between age groups, semantic tasks and their interaction for each pair of seed ROIs.

#### Whole-brain-level analyses

Here, we explored how the core semantic areas interacted with other brain regions during semantic cognition. For each participant and each ROI seed, we used the PPI term effect maps which contrasted each semantic task with the non-semantic task (i.e., semantic knowledge vs. non-semantic, semantic control vs. non-semantic) and between the two semantic tasks (i.e., knowledge vs. control) as the first-level input. We submitted the first-level contrast maps to second-level analyses for group effects. First, by conducting one-sample t-tests over the semantic knowledge/control vs. non-semantic maps across all participants, we tested where connectivity with the seeds varied between each type of semantic task and non-semantic processing. By performing two-sample t-tests, we examined the whole-brain-level connectivity differences between older and young people during the semantic tasks relative to the non-semantic task. Second, we examined connectivity differences between semantic control and semantic knowledge tasks. One-sample t-tests across all participants were used to test the overall differences in PPI patterns between semantic control versus knowledge tasks, and two-sample t-tests were used to compare the age group differences in this contrast.

## Results

### Univariate activation analysis

Before the investigation of functional connectivity, we first conducted a univariate analysis to identify brain regions that were activated during each semantic task, compared with a baseline of non-semantic executive processing. As reported by Wu and Hoffman (2023), for the semantic control task, positive effects were found in core semantic regions including IFG and ATL, as well as DMN regions including angular gyrus, posterior cingulate and ventromedial prefrontal cortex (Figure 2; for activation peaks, see Supplementary Table 1). In contrast, higher activation for the non-semantic task was found in a wide range of areas across parietal and frontal lobes, which overlap with MDN. Older people activated DMN regions more than the young, while young adults showed more activation in the MDN regions of the insula, presupplementary motor area and posterior inferior temporal gyrus. For the semantic knowledge task, similar sets of regions were found for both the overall effects and age group differences, although the IFGs’ preference for semantic processing became less prominent here.

**Figure 2.**
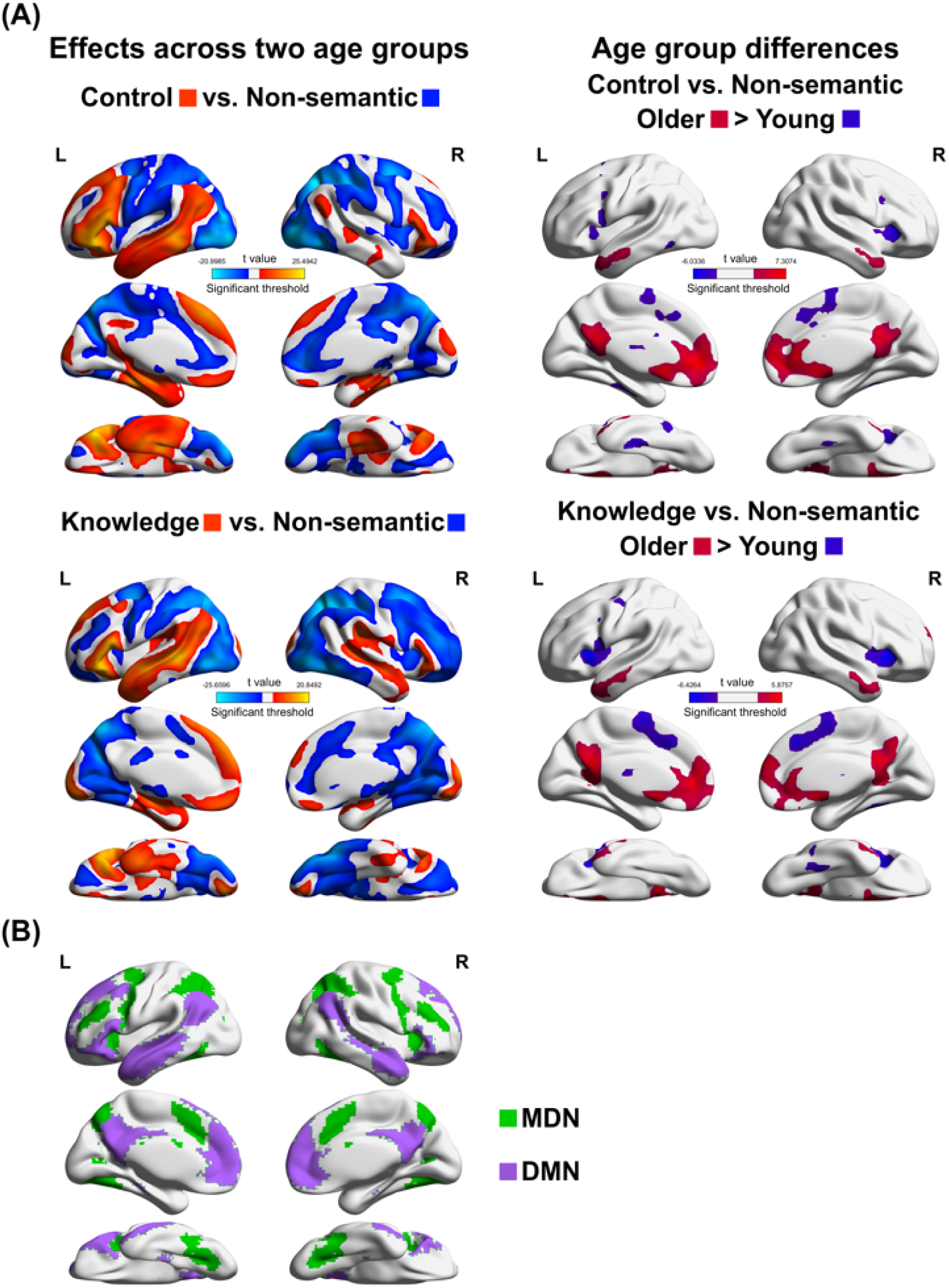
Results of whole-brain univariate activation analysis. (A) shows the univariate activation effects across the two age-groups for the contrasts of each semantic task versus the non-semantic task (left), and the age group differences (right). For comparison, (B) shows large-scale brain networks MDN (Fedorenko, Duncan, & Kanwisher, 2013) and DMN (Yeo et al., 2011), as identified in previous studies. Results were corrected for multiple comparisons, voxelwise p < 0.005, FWE corrected cluster threshold p = 0.05.

### ROI-level functional connectivity

To examine how semantic processing modulated the connectivity between core semantic regions, we performed a set of ROI-level analyses. Specifically, gPPI was used to test how functional connectivity between left and right vATLs and IFGs changed when participants engaged in semantic processing (with the non-semantic task as the baseline). As shown in Table 1 and Figure 3, across all participants, the left IFG exhibited overall increased connectivity with the right IFG and left vATL during the semantic tasks (both *t* >= 4.75, both two-tailed *p* < 10^-4^). Thus, semantic processing elicited increased interaction between the core semantic control area (i.e., left IFG) and both the knowledge area (i.e., left vATL) and the semantic control area in the contralateral hemisphere (i.e., right IFG). The remaining connections showed effects in the same direction but these were not statistically significant (all *t* _169_ <= 1.77, all two-tailed *p* >= 0.12).

**Table 1.**
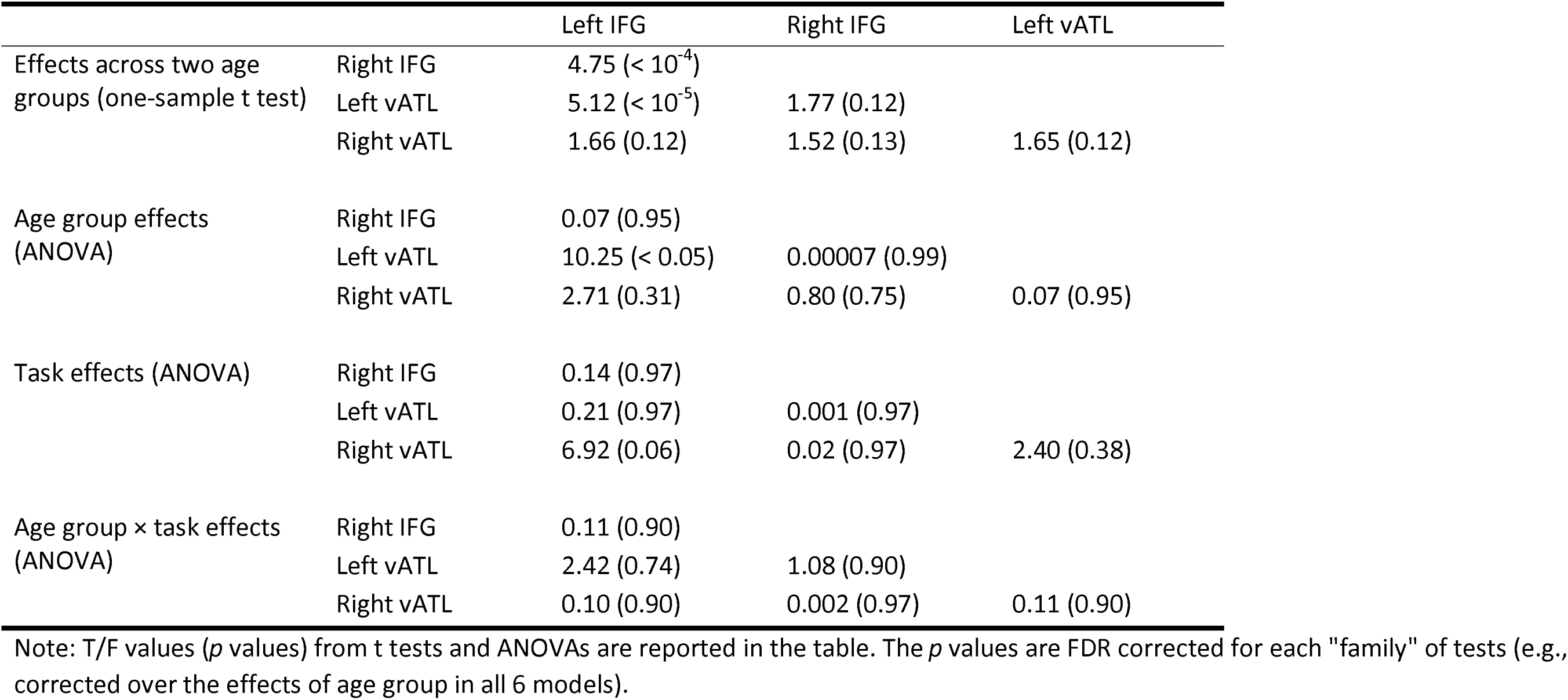
Effects of the ROI-level PPI analysis for the semantic tasks (vs. the non-semantic task)

|  |  | Left IFG | Right IFG | Left vATL |
| --- | --- | --- | --- | --- |
| Effects across two age groups (one-sample t test) | Right IFG | 4.75 ($< 10^{-4}$) | | |
| | Left vATL | 5.12 ($< 10^{-5}$) | 1.77 (0.12) | |
|  | Right vATL | 1.66 (0.12) | 1.52 (0.13) | 1.65 (0.12) |
| Age group effects (ANOVA) | Right IFG | 0.07 (0.95) |  |  |
| | Left vATL | 10.25 ($< 0.05$) | 0.00007 (0.99) | |
|  | Right vATL | 2.71 (0.31) | 0.80 (0.75) | 0.07 (0.95) |
| Task effects (ANOVA) | Right IFG | 0.14 (0.97) |  |  |
|  | Left vATL | 0.21 (0.97) | 0.001 (0.97) |  |
|  | Right vATL | 6.92 (0.06) | 0.02 (0.97) | 2.40 (0.38) |
| Age group × task effects (ANOVA) | Right IFG | 0.11 (0.90) | | |
|  | Left vATL | 2.42 (0.74) | 1.08 (0.90) |  |
|  | Right vATL | 0.10 (0.90) | 0.002 (0.97) | 0.11 (0.90) |

Note: T/F values ( $p$ values) from t tests and ANOVAs are reported in the table. The $p$ values are FDR corrected for each "family" of tests (e.g., corrected over the effects of age group in all 6 models).

**Figure 3.**
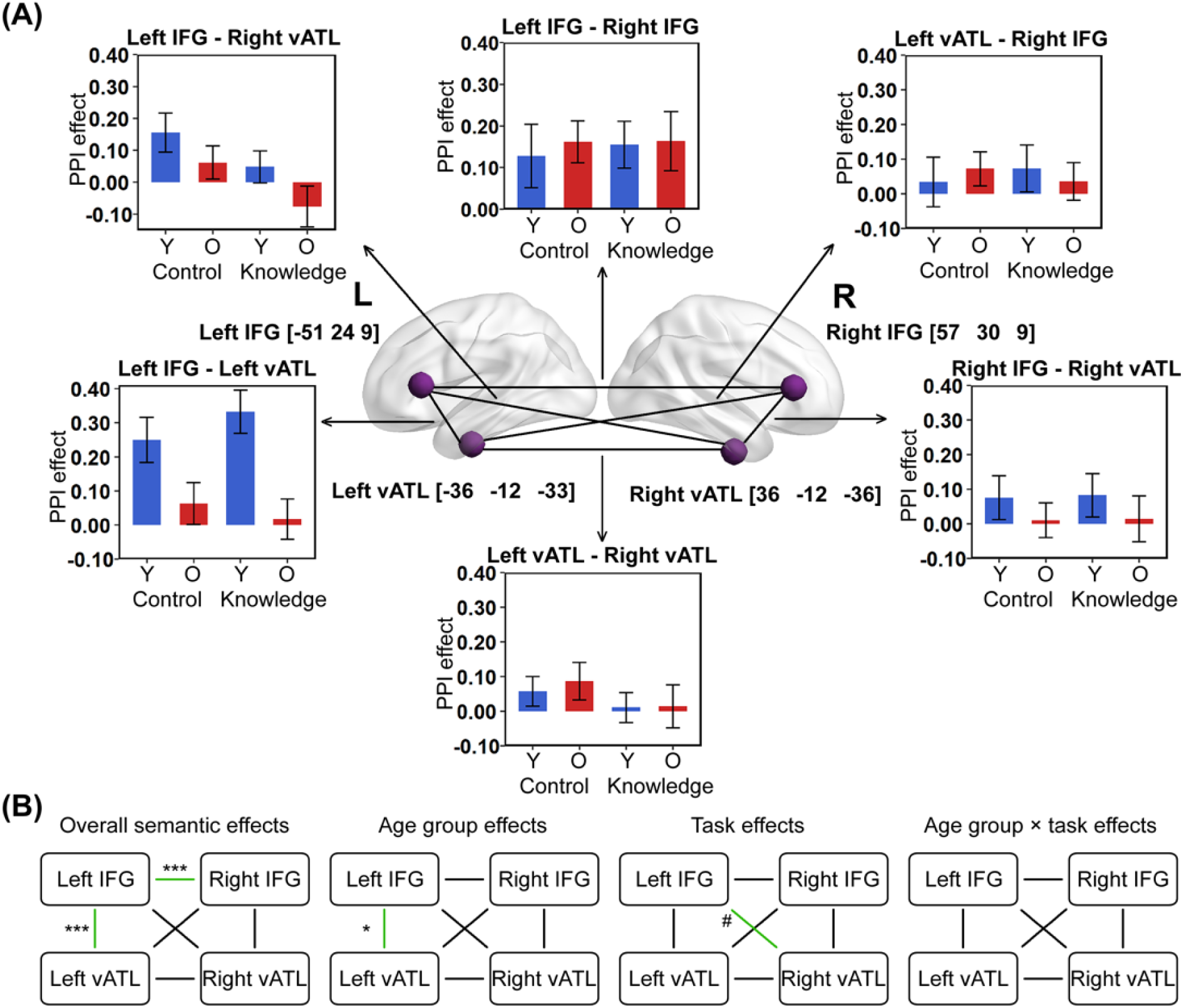
Group-level seed coordinates and results of the ROI-level PPI analysis. (A) shows the PPI effects in two age groups and two semantic tasks with the non-semantic task as the baseline Error bars indicate SEM. (B) indicates which connections showed significant effects of the experimental manipulations. Asterisks indicate significance levels (FDR-corrected for each “family” of tests), * p < 0.05, ** p < 0.01, *** p < 0.001, # marginally significant p = 0.06.

We then performed follow-up ANOVAs to further investigate if age group and type of semantic task affected connectivity between semantic regions (Table 1 and Figure 3). There were significant age group differences in the connectivity between the left IFG and left vATL for semantic processing (*F* _(1,83)_ = 10.25, *p* < 0.05). In the post-hoc two-sample t-test, we found this age group effect reflected a greater connectivity increases for semantic processing in the young group (*t* _168_ = 4.02, two-tailed *p* < 10^-4^). The ANOVAs also showed a marginally significant task effect for the connectivity between the left IFG and right vATL (*F* _(1,83)_ = 6.92, *p* = 0.06), which indicated a stronger connectivity between the left IFG and right vATL during the semantic control task compared with the knowledge task (post-hoc paired-sample t test, *t* _84_ = 2.64, *p* < 0.01). No interactions between age group and type of semantic task were found (all *F* _(1,83)_ <= 2.42, all *p* >= 0.74).

### Whole-brain-level functional connectivity

In this section, we explored areas across the whole brain whose connectivity with the core semantic areas was modulated by semantic processing, and how healthy ageing affects this connectivity. We started by examining the overall PPI effects of the semantic control and knowledge tasks (relative to the non-semantic task) across all participants (Figures 4 and 5; for peak effect co-ordinates, see Supplementary Tables 2 and 3). Connectivity effects were strongest for the left IFG seed and weakest for the right vATL seed, which is consistent with this region’s lower level of activation in the semantic tasks (Figure 2). In general, however, a consistent set of regions showed increased connectivity with the four seed ROIs during semantic processing, irrespective of semantic task. These regions included occipital cortices and MDN areas such as the intraparietal sulcus (IPS), posterior inferior temporal gyrus, dorsolateral prefrontal cortex and supplementary motor areas. This increased coupling occurred despite the fact that many of the same areas showed the opposite pattern in activation, i.e., lower activity during semantic processing (see Figure 2 and Wu and Hoffman, 2023). Conversely, reduced connectivity with the seed regions was found mostly in nodes of the DMN, including angular gyrus, posterior cingulate, and medial frontal cortex. Counter-intuitively, these regions tended to show increased activity during semantic tasks cf. non-semantic processing (Figure 2). Thus, at this macro scale, the overall picture is that activation and functional connectivity do not go hand in hand.

**Figure 4.**
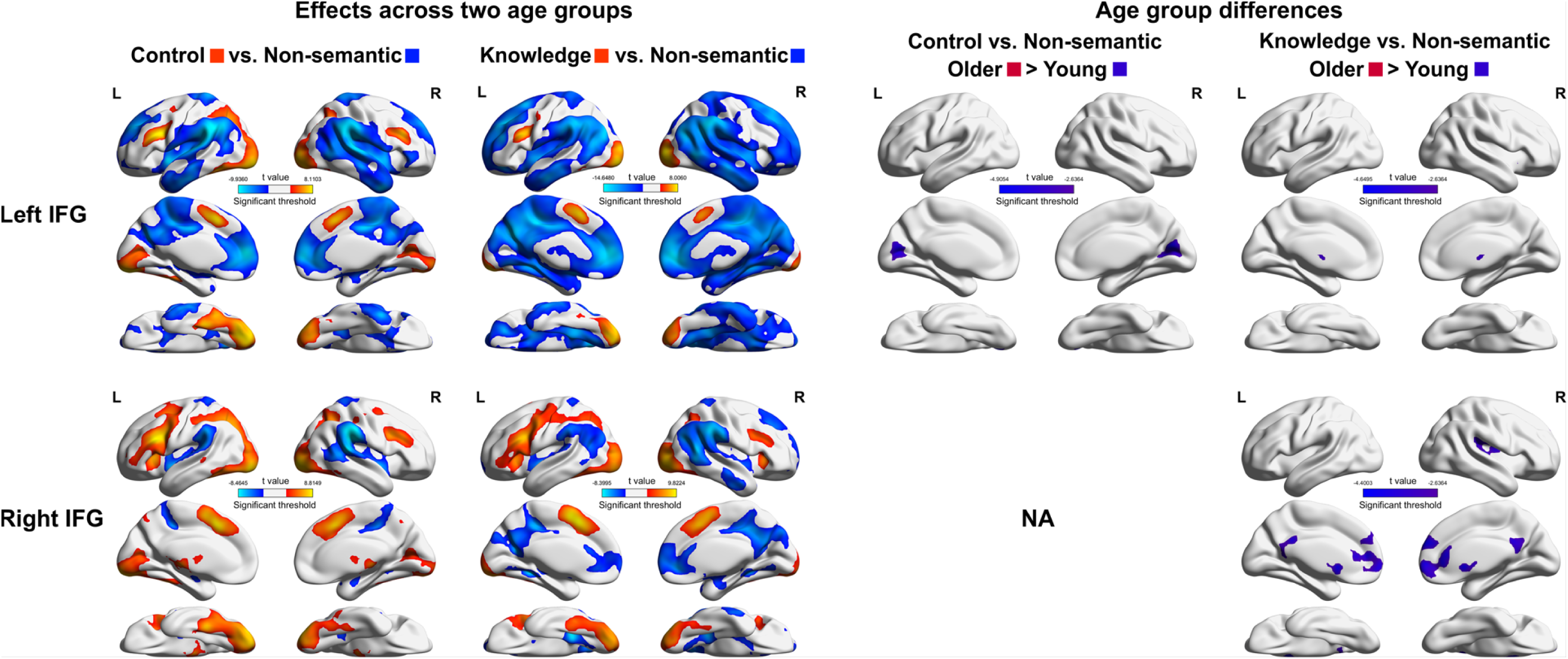
Results of the whole-brain PPI analysis with left IFG and right IFG as seeds. This figure shows the PPI effect maps across two age groups (the first two columns), for each of the semantic tasks contrasted with the non-semantic task. We also compared the PPI effects between older and young people (the last two columns). In the all-participants effect maps, hot colours indicate semantic task PPI effects and cold colours indicate non-semantic task PPI effects. In the age-group-differences maps, hot colours indicate older adults had stronger PPI effects and cold colours indicate greater PPI effects in the young. Result maps were corrected for multiple comparisons, voxelwise p < 0.005, FWE corrected cluster threshold p = 0.05.

**Figure 5.**
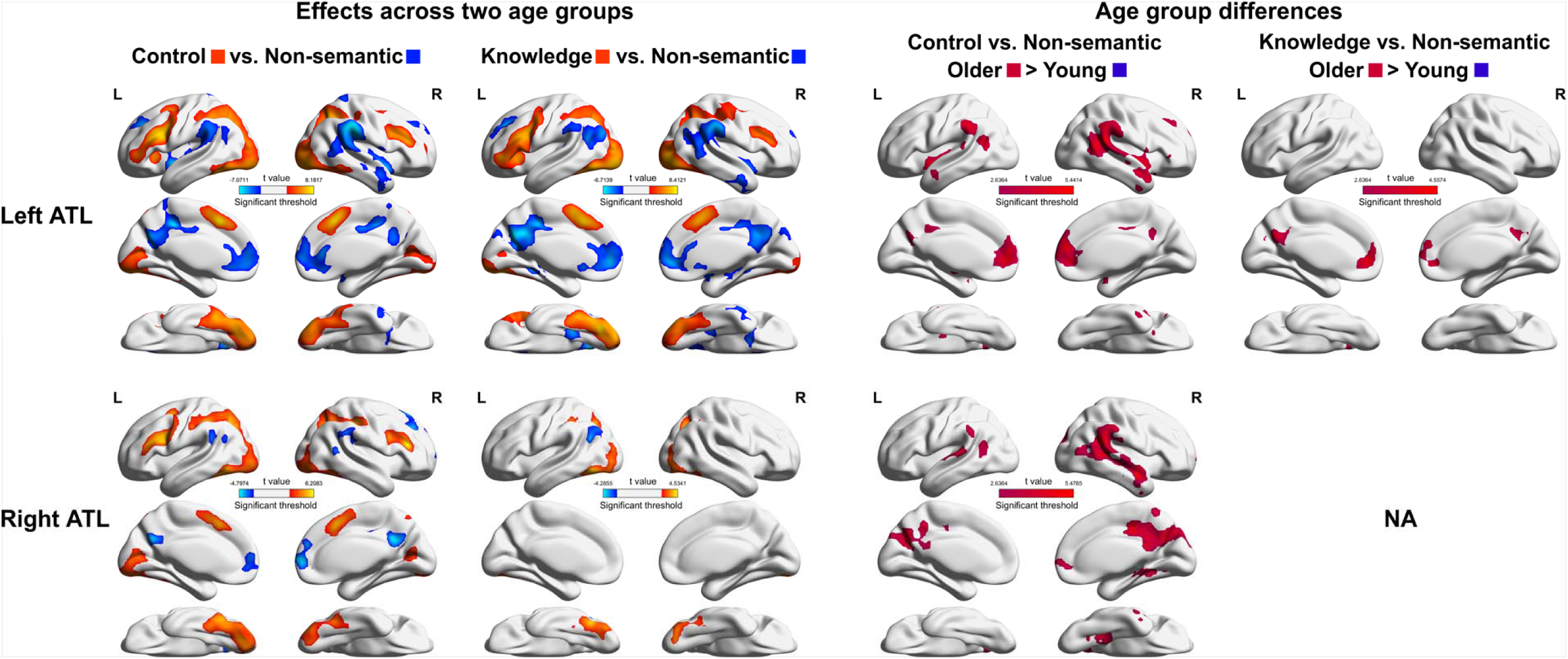
Results of the whole-brain PPI analysis with left ATL and right ATL as seeds. This figure shows the PPI effect maps across two age groups (the first two columns), for each of the semantic tasks contrasted with the non-semantic task. We also compared the PPI effects between older and young people (the last two columns). In the all-participants effect maps, hot colours indicate semantic task PPI effects and cold colours indicate non-semantic task PPI effects. In the age-group-differences maps, hot colours indicate older adults had stronger PPI effects and cold colours indicate greater PPI effects in the young. Result maps were corrected for multiple comparisons, voxelwise p < 0.005, FWE corrected cluster threshold p = 0.05.

We also evaluated age group differences for the above effects with two-sample t-tests. When left IFG was the seed, young adults exhibited greater connectivity between left IFG and early visual areas in the semantic control task, and more connectivity between left IFG and caudate for the semantic knowledge task. When right IFG was the seed, young adults’ right IFG showed more coupling with midline DMN regions in the knowledge task compared with older people. The left vATL seed had more interaction with the DMN regions in older adults in the two semantic tasks. A similar age group difference was also found for the right vATL seed, but only for the semantic control task.

In the second part of the whole-brain-level PPI analysis, we compared the PPI effects in the two semantic tasks. As shown in Figure 6, the seeds of left IFG and right IFG connected more with a wide array of regions during the semantic control task than the knowledge task (for peak effect co-ordinates, see Supplementary Table 4). A comparison between age groups revealed that older adults’ left IFG was more connected with the medial prefrontal cortex during the semantic control task, while young people’s left IFG was more connected with the calcarine/lingual areas during the control task. Weaker effects were found for the seeds of left and right vATLs. Specifically, the left vATL connected more with the precuneus during the semantic control task cf. the knowledge task. The right vATL showed an age group difference, in which older people’s right vATL was more connected with cuneus and posterior cingulum in the semantic control task than the young.

**Figure 6.**
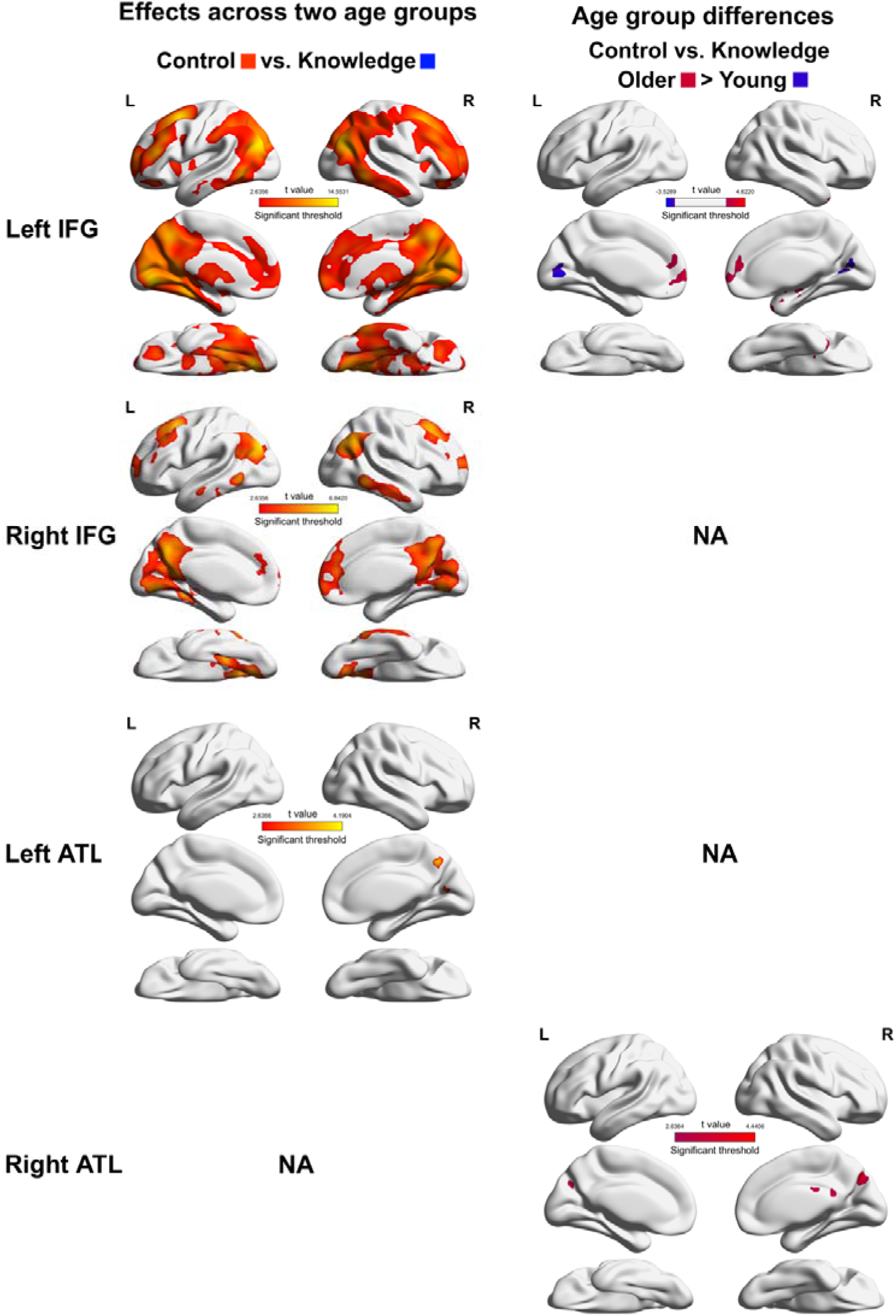
Results of the whole-brain PPI analysis for semantic task effects, with each of the semantic ROIs as the seed. This figure shows the PPI effect maps across two age groups for the two semantic tasks contrasting with each other (left). We also compared the PPI effects between older and young people (right). In the all-participants effect maps, hot colours indicate semantic control task PPI effects and cold colours indicate semantic knowledge task PPI effects. In the age- group-differences maps, hot colours indicate older adults had stronger PPI effects and cold colours indicate greater PPI effects in the young. Result maps were corrected for multiple comparisons, voxelwise p < 0.005, FWE corrected cluster threshold p = 0.05.

## Discussion

To further our understanding of the neural dynamics supporting semantic cognition, the current study systematically investigated connectivity of the core semantic network (i.e., left and right IFGs and vATLs) during two semantic tasks in older and young adults. Left IFG, a critical region for control over semantic processing, exhibited more collaboration with the right IFG and left vATL during the semantic tasks, when compared with a non-semantic task. With respect to ageing effects, connectivity between left IFG and vATL during semantic processing was stronger in young people than in the old. At a whole-brain level, all four semantic regions demonstrated a largely similar whole-brain connectivity pattern. Semantic tasks induced more co-operation between semantic regions and domain-general executive control areas (i.e., the MDN) and more decoupling between semantic regions and parts of the DMN. These results ran counter to task-related activation in these networks, as DMN regions generally showed activation increases in response to semantic tasks while MDN showed decreases. These findings have implications for understanding functional interactions in the semantic network and for understanding the participation of large-scale brain networks in semantic cognition in different age groups.

### Interactions between IFGs and vATLs

Our study reveals the dynamic interaction between the left IFG and other semantic areas. Aligning with our expectations, we found that semantic processing increased left-right IFG connectivity. The role of right IFG in language processing has long been a matter of debate. Small amounts of right IFG activation during language tasks have been interpreted either as a compensatory response to high processing demands or as dedifferentiated activation of functionally-irrelevant cortex (Gainotti, 2015; Hoffman & Morcom, 2018; Jung et al., 2021; Krieger-Redwood et al., 2015; Quillen et al., 2021; Stefaniak et al., 2020; Turkeltaub, 2015). In a recent study, we found that both left and right IFG activation increased linearly with task demands in different semantic tasks and across young and older age groups (Wu & Hoffman, 2023). This result suggests that right IFG activation is functionally significant for semantic tasks, consistent with a supporting role in semantic cognition. In the current study, the increased connectivity between left and right IFGs provides a new source of evidence for this view. Together, both our previous activation results and the current connectivity results suggest that the right IFG collaborates with the left to support semantic cognition.

We also found increased connectivity between the left IFG and left vATL during semantic tasks, consistent with previous research (Jung et al., 2021). According to the CSC framework (Jefferies, 2013; Lambon Ralph et al., 2017), IFG supports semantic cognition by regulating the activation and use of knowledge representations coded primarily in the ATLs. Our results support this view and provide evidence for interaction between knowledge and control resources to achieve goal-directed semantic processing. Strengthened knowledge-control coupling was found for the left IFG – left vATL connection, but not for the parallel connections involving right IFG (i.e., right IFG – left vATL and right IFG – right vATL). Connectivity effects involving right IFG were positive for semantic tasks but not statistically significant, so it is possible that these effects were simply weaker and our data do not have sufficient power to detect them. This is understandable given the evidence that right IFG plays a less central role in semantic processing.

The left IFG – right vATL connection was also not significantly enhanced by the semantic tasks. One potential reason for this non-significant result is that right vATL was less centrally involved in the tasks used here. Although the bilateral vATLs have together been implicated in semantic knowledge representation, they do show graded specialisation in function with evidence for a left-lateralized bias for written word semantic processing (Gainotti, 2012, 2014; Hoffman & Lambon Ralph, 2018; Rice, Hoffman, et al., 2015; Rice, Lambon Ralph, et al., 2015; Snowden et al., 2004). The written word stimuli in the current study do not appear to have engaged right vATL as strongly as the left, which may explain why this region did not show increased connectivity with left-hemisphere semantic regions. This argument is supported by the significant task effect we found in the connection between left IFG and right vATL. Our results showed an enhanced collaboration between the left IFG and right vATL in the semantic control task compared to the knowledge task. Though both used written words, the semantic control task required participants to make judgements about non-verbal semantic properties (typical colour or size of objects). Non-verbal semantic processing elicits more bilateral vATL activity (Rice, Lambon Ralph, et al., 2015), which might lead to more interactions involving right vATL in this task.

We also investigated healthy ageing’s effect on dynamics of the semantic network. The DECHA framework has described a shifting reliance with age from fluid intelligence (i.e., executive control abilities) to crystallized intelligence (i.e., experience and knowledge) and links this shifting cognitive architecture to less flexible connectivity between control and knowledge-supporting regions in the ageing brain (Spreng & Turner, 2019; Turner & Spreng, 2015). Previous studies which tested the DECHA predictions have either focused on non-semantic tasks or have looked at connectivity between large-scale networks (Adnan, Beaty, Silvia, Spreng, & Turner, 2019; Martin, Saur, & Hartwigsen, 2022; Martin, Williams, Saur, & Hartwigsen, 2023; Spreng et al., 2018; Wu & Hoffman, 2023). The current study tested the DECHA prediction more precisely within the core semantic regions. We found that the young participants showed a larger left IFG – left ATL connectivity difference for semantic versus non-semantic tasks than the old. This finding suggests that young people show more flexible connectivity between knowledge and control-supporting parts of the semantic network, adjusting the interaction between these regions depending on the relevance of semantics to the task. This supports the DECHA connectivity prediction in the domain of semantic cognition. Our results also suggest that future studies examining DECHA, in addition to considering broad neural networks, also investigate the specific regions that are contributing relevant knowledge and control processes to the task under study.

A related potential explanation for our age group effect is that it reflects specific differences in semantic abilities between age groups. Young people have less developed semantic knowledge than the old (Hoffman, 2018; Salthouse, 2004; Verhaeghen, 2003), so may be more reliant on control processes to search and shape their existing knowledge and retrieve task-relevant information (Hoffman, 2018; Spreng & Turner, 2019; Turner & Spreng, 2015). This explanation has parallels with data from neuropsychological studies, where patients with impaired semantic knowledge due to ATL damage or resection showed increased IFG activation (Billingsley, McAndrews, Crawley, & Mikulis, 2001; Rice et al., 2018). This explanation is also compatible with the DECHA account, as both emphasize age-related differences in the interaction between knowledge and control resources.

### Connectivity between core semantic regions and other networks

The present study also explored how each of the core semantic regions connected with the other regions during semantic processing. We found frequent convergence across the whole-brain connectivity patterns for the two semantic tasks (i.e., knowledge and control tasks) and the four ROIs (i.e., left and right IFGs and vATLs). Specifically, the semantic tasks increased the connectivity between the core semantic regions and MDN domain-general executive control regions (e.g., IPS and supplementary motor areas) and parts of visual cortex. In contrast, the semantic ROIs decoupled from DMN regions during semantic tasks, including angular gyrus, posterior cingulate, and medial frontal cortex.

The engagement of executive networks for language and semantic processing is a current area of debate. The CSC framework proposes specialised control regions for semantic processing that have limited overlap with the domain-general control regions of the MDN (for example see Jackson, 2021). Other researchers have also argued that language processing does not engage domain-general control regions and that cognitive control for language is instead supported by specialised regions (Diachek, Blank, Siegelman, Affourtit, & Fedorenko, 2020; Fedorenko & Thompson-Schill, 2014; Quillen et al., 2021). Some aspects of the current data support this view: we found that MDN areas were less active during semantic tasks than during a cognitively demanding non-semantic task. However, other aspects of the data suggest active involvement of MDN in semantic processing. First, in earlier analyses, we found that MDN activity increased with task demands for both semantic and non-semantic tasks (though the effect was stronger for the non-semantic task) (Wu & Hoffman, 2023). Second, our present results show that the MDN increases connectivity with core semantic areas during the performance of semantic tasks. Overall, this pattern of results indicates that MDN regions actively contribute to semantic language tasks, albeit to a lesser extent than for other types of cognitive demand. Our findings also indicate that we should be cautious about relying only on subtraction analyses to make inferences about function of these regions. Less engagement in semantic tasks does not appear to be synonymous with no engagement at all.

We also found paradoxical effects in the connectivity between core semantic regions and DMN regions. Although DMN regions were more active for semantic than non-semantic tasks, their connectivity with IFG and vATL decreased during semantic processing. DMN regions support a range of memory and experience-driven processes, including integrative semantic processing, recollections of episodic memories, social cognition, and constructing models of ongoing events and situations (Binder & Desai, 2011; Smallwood, Bernhardt, et al., 2021; Smallwood, Turnbull, et al., 2021; Speer, Zacks, & Reynolds, 2007; Yeshurun, Nguyen, & Hasson, 2021). In semantic tasks, stimulus-driven DMN activity has been associated with automatic retrieval of associated experiences and knowledge (Wang, Margulies, Smallwood, & Jefferies, 2020). This can explain why our semantic trials engaged DMN more than the non-semantic task: meaningful concepts automatically activate representations of a range of associated memories, events and contexts, in a way that meaningless letter strings do not. However, our semantic tasks required specific, decontextualised judgements about particular properties or linguistic associations, so this broader knowledge about the situational, social or episodic aspects of concepts was not relevant. Accordingly, the core semantic network decoupled from DMN regions during task performance, in order to prioritise the specific lexical and visual properties required by the tasks. This explanation is supported by our previous finding that DMN activity was not correlated with semantic task demands in the present data (Wu & Hoffman, 2023).

Our account proposes that DMN regions will tend to be automatically activated by meaningful stimuli (relative to less meaningful stimuli), but that the functional roles of these regions depends on specific task demands. Integrative or contextualised semantic processing is likely to rely on DMN regions to a greater extent. Indeed, previous research using verbal fluency (Martin et al., 2022; Martin et al., 2023) and creative thinking tasks (Adnan et al., 2019) has found enhanced DMN connectivity which is correlated with better task performance, at least in young adults. In these tasks, activating the broader situational aspects of concepts is helpful (for example, when generating exemplars of animals, it is useful to imagine contexts like farms and zoos). In our study, in contrast, IFG and ATL are co-activated with DMN regions but they do not seem to be acting as a single functional network.

Additionally, our results revealed an age-related effect in the connectivity between bilateral vATLs and DMN regions: the decoupling of DMN regions from vATL during semantic tasks was less in older people than the young. This effect may indicate that older people’s semantic-specific knowledge is more entangled with their broader experiences and episodic memories, making it more difficult for the older group to separate the corresponding neural correlates. In contrast, older people showed more decoupling between the right IFG and DMN midline areas in the knowledge task. Since the representations coded in these DMN regions were not relevant for our tasks, their decoupling from IFGs might indicate inhibition of these irrelevant associations. The stronger effect in older people could indicate that right IFG is more involved in these regulatory processes in later life. Finally, young people’s left IFG was more strongly connected with visual cortex than older adults during the control task. According to the CSC framework, sensory-motor regions in the brain work as functionally diverse ‘spokes’ to represent modality-specific knowledge (Lambon Ralph et al., 2017). The enhanced connectivity between left IFG and visual cortex may indicate that, during semantic judgements that involve more embodied visual information (i.e., colour/size pairing in the control task), young people are better able to enhance communication between semantic control regions and relevant spoke areas to facilitate task processing.

In conclusion, focusing on the core semantic regions of bilateral IFGs and vATLs, our study has provided a comprehensive view on dynamic interactions within the semantic network, as well as between these areas and the rest of the brain. We found strong interactions between the left IFG and the other semantic regions during semantic tasks, with additional effects of cognitive aging and type of semantic task. At a whole-brain level, semantic tasks in the current study involved interactions between core semantic regions and some domain-general executive areas outside the semantic network. DMN regions, on other hand, were activated by meaningful stimuli but did not appear to collaborate with core semantic areas. Together, our findings paint a complex picture of cooperation for semantic cognition within and beyond the core semantic network.

## Acknowledgements

This work was supported by a BBSRC grant to P.H. (BB/T004444/1). Imaging was carried out at the Edinburgh Imaging Facility (www.ed.ac.uk/edinburgh-imaging), University of Edinburgh, which is part of the SINAPSE collaboration (www.sinapse.ac.uk). We are grateful to the University of Minnesota Center for Magnetic Resonance Research for sharing their neuroimaging sequences. We thank Yueyang Zhang for his help with data collection. We are also grateful to all research participants in this study. For the purpose of open access, the authors have applied a Creative Commons Attribution (CC BY) licence to any Author Accepted Manuscript version arising from this submission.

## CRediT authorship contribution statement

Wei Wu: Conceptualization, Investigation, Formal analysis, Validation, Writing - original draft, Writing - review & editing, Visualization. Paul Hoffman: Conceptualization, Writing - review & editing, Project administration, Supervision, Funding acquisition.

