## Supplementary Table for "Functional integration and segregation during semantic cognition: Evidence across age groups"

**Supplementary Information**

**Overview**

| Supplementary Table S1 | Peak MNI coordinates for the results of whole-brain univariate activation analysis |
| --- | --- |
| Supplementary Table S2 | Peak MNI coordinates for the results of the whole-brain PPI analysis with left IFG and right IFG as seeds |
| Supplementary Table S3 | Peak MNI coordinates for the results of the whole-brain PPI analysis with left ATL and right ATL as seeds |
| Supplementary Table S4 | Peak MNI coordinates for the results of the whole-brain PPI analysis for semantic task effects, with each of the ROIs as the seed. |

**Table S1. Peak MNI coordinates for the results of whole-brain univariate activation analysis.**

|  | Effects across two age groups | | | | |  |  | Age group differences | | | | |
| --- | --- | --- | --- | --- | --- | --- | --- | --- | --- | --- | --- | --- |
|  | Region | x | y | z | t value |  |  | Region | x | y | z | t value |
| Control  vs.  Non-semantic | L IFG (pars orbitalis) | -42 | 36 | -12 | 25.49 |  | Control  vs.  Non-semantic  Older > Young | L MPFC | -9 | 48 | 0 | 7.31 |
|  | L IFG (pars triangularis) | -48 | 24 | 9 | 19.80 |  |  | Olfactory | 0 | 15 | -9 | 7.16 |
|  | L MTG | -54 | -42 | -6 | 18.71 |  |  | R PCC | 3 | -45 | 27 | 6.46 |
|  | L MPFC | -9 | 39 | 42 | 17.02 |  |  | R Linual Gyrus | 9 | -45 | 6 | 3.78 |
|  | L Fusiform Gyrus | -30 | -30 | -21 | 16.43 |  |  | L Temporal Pole | -51 | 15 | -30 | 4.92 |
|  | L Fusiform Gyrus | -33 | -9 | -33 | 14.16 |  |  | L MTG | -66 | -6 | -15 | 4.90 |
|  | R Cerebelum | 30 | -72 | -42 | 22.26 |  |  | R MTG | 66 | -6 | -15 | 4.40 |
|  | R Cerebelum | 15 | -84 | -36 | 20.45 |  |  | R Temporal Pole | 51 | 9 | -27 | 4.21 |
|  | L Cerebelum | -15 | -84 | -36 | 7.83 |  |  | L Cerebelum | -42 | -60 | -30 | -6.03 |
|  | R IFG (parts orbitalis) | 45 | 36 | -15 | 13.88 |  |  | R Cerebelum | 36 | -48 | -30 | -5.28 |
|  | R PHG | 33 | -27 | -21 | 11.91 |  |  | Cerebelum | 0 | -60 | -21 | -5.01 |
|  | R IFG (pars triangularis) | 57 | 30 | 12 | 9.92 |  |  | L Putamen | -21 | -3 | 9 | -4.68 |
|  | R Fusiform Gyrus | 33 | -12 | -36 | 9.65 |  |  | L MPFC | 0 | 27 | 45 | -4.44 |
|  | R MTG | 57 | -69 | 21 | 7.55 |  |  | R Insula | 33 | 24 | 3 | -4.71 |
|  | R MTG | 51 | -36 | -3 | 5.39 |  |  | R Putamen | 30 | -12 | 3 | -4.03 |
|  | L Occipital | -9 | -102 | 12 | 9.42 |  |  | R IFG (pars triangularis) | 45 | 15 | 30 | -3.11 |
|  | R Calcarine Gyrus | 12 | -96 | 18 | 8.36 |  |  |  |  |  |  |  |
|  | L Cerebelum | -18 | -78 | -9 | 4.65 |  |  |  |  |  |  |  |
|  | R Precuneus | 21 | -69 | 45 | -21.00 |  |  |  |  |  |  |  |
|  | R Occipital | 36 | -87 | -3 | -19.90 |  |  |  |  |  |  |  |
|  | L Occipital | -30 | -87 | 15 | -19.38 |  |  |  |  |  |  |  |
|  | L Occipital | -30 | -93 | -6 | -19.20 |  |  |  |  |  |  |  |
|  | L SPL | -15 | -69 | 48 | -17.16 |  |  |  |  |  |  |  |
|  | R Occipital | 42 | -66 | -9 | -16.76 |  |  |  |  |  |  |  |
|  | R Supramarginal Gyrus | 39 | -39 | 48 | -16.74 |  |  |  |  |  |  |  |
|  | R SFG | 21 | 3 | 57 | -16.56 |  |  |  |  |  |  |  |
|  | L SFG | -24 | -3 | 51 | -14.43 |  |  |  |  |  |  |  |
|  | L IPL | -30 | -42 | 48 | -14.01 |  |  |  |  |  |  |  |
|  | R IFG (pars opercularis) | 48 | 3 | 27 | -13.37 |  |  |  |  |  |  |  |
|  | R MCC | 12 | -39 | 45 | -13.09 |  |  |  |  |  |  |  |
| Knowledge  vs.  Non-semantic | R Cerebelum | 27 | -75 | -39 | 20.85 |  | Knowledge  vs.  Non-semantic  Older > Young | L Precuneus | 0 | -63 | 33 | 5.88 |
|  | L Occipital | -12 | -102 | 6 | 16.21 |  |  | R Linual Gyrus | 15 | -51 | 6 | 5.59 |
|  | R Cuneus | 15 | -102 | 12 | 15.05 |  |  | L Linual Gyrus | -12 | -51 | 6 | 5.25 |
|  | L IFG (pars orbitalis) | -51 | 27 | -3 | 19.05 |  |  | L MPFC | 0 | 54 | 9 | 5.36 |
|  | L MTG | -57 | -42 | 0 | 17.48 |  |  | R MPFC | 0 | 30 | -6 | 4.88 |
|  | L MTG | -54 | -9 | -9 | 13.25 |  |  | R Temporal Pole | 51 | 15 | -30 | 4.30 |
|  | L SFG | -9 | 48 | 39 | 14.33 |  |  | L MTG | -60 | -9 | -18 | 3.91 |
|  | L SMA | -6 | 18 | 60 | 11.20 |  |  | L Insula | -33 | 15 | -3 | -6.43 |
|  | L Rectal Gyrus | -3 | 48 | -18 | 9.43 |  |  | L Caudate | -9 | 6 | 18 | -5.99 |
|  | R Temporal Pole | 42 | 6 | -21 | 10.97 |  |  | R Insula | 36 | 21 | 0 | -6.14 |
|  | R STG | 51 | -30 | 21 | 10.49 |  |  | R Putamen | 30 | -12 | 3 | -4.82 |
|  | R MTG | 48 | -33 | 0 | 9.79 |  |  | L Cerebelum | -42 | -60 | -30 | -5.61 |
|  | R AG | 27 | -63 | 48 | -25.66 |  |  | Cerebellar Vermis | -3 | -63 | -33 | -4.78 |
|  | R Occipital | 30 | -72 | 30 | -23.98 |  |  | R MCC | 6 | 30 | 42 | -5.37 |
|  | L Occipital | -18 | -69 | 30 | -22.50 |  |  | L MCC | -6 | 9 | 45 | -4.64 |
|  | L SPL | -21 | -66 | 51 | -22.13 |  |  | R Cerebelum | 33 | -48 | -30 | -4.64 |
|  | L IPL | -45 | -36 | 42 | -21.70 |  |  | R ITG | 54 | -60 | -18 | -3.71 |
|  | R Precuneus | 12 | -75 | 54 | -20.53 |  |  |  |  |  |  |  |
|  | R Supramarginal Gyrus | 42 | -39 | 48 | -20.49 |  |  |  |  |  |  |  |
|  | R Occipital | 42 | -78 | -3 | -20.36 |  |  |  |  |  |  |  |
|  | R ITG | 51 | -54 | -12 | -20.14 |  |  |  |  |  |  |  |
|  | L Occipital | -45 | -72 | -6 | -19.58 |  |  |  |  |  |  |  |
|  | R MFG | 24 | 12 | 54 | -19.19 |  |  |  |  |  |  |  |
|  | L MFG | -24 | 0 | 54 | -17.53 |  |  |  |  |  |  |  |

Note: AG, angular gyrus; IFG, inferior frontal gyrus; IPL, inferior parietal lobule; ITG, inferior temporal gyrus; MCC, middle cingulate cortex; MFG, middle frontal gyrus; MPFC, medial prefrontal cortex; MTG, middle temporal gyrus; PCC, posterior cingulate cortex; PHG, parahippocampal gyrus; SMA, supplementary motor area; SFG, superior frontal gyrus; SPL, superior parietal lobule; STG, superior temporal gyrus.

**Table S2. Peak MNI coordinates for the results of the whole-brain PPI analysis with left IFG and right IFG as seeds.**

|  | Effects across two age groups | | | | |  |  | Age group differences | | | | |
| --- | --- | --- | --- | --- | --- | --- | --- | --- | --- | --- | --- | --- |
|  | Region | x | y | z | t value |  |  | Region | x | y | z | t value |
| Seed left IFG  Control  vs.  Non-semantic | L IFG (pars triangularis) | -42 | 15 | 24 | 8.11 |  | Seed left IFG  Control  vs.  Non-semantic  Older > Young | R Calcarine Gyrus | 6 | -69 | 15 | -4.91 |
|  | L Precentral Gyrus | -30 | 0 | 51 | 4.72 |  |  | R Fusiform Gyrus | 24 | -78 | -3 | -3.09 |
|  | L Fusiform Gyrus | -24 | -84 | -12 | 7.85 |  |  |  |  |  |  |  |
|  | L Fusiform Gyrus | -36 | -39 | -21 | 7.30 |  |  |  |  |  |  |  |
|  | R Linual Gyrus | 21 | -90 | -9 | 7.19 |  |  |  |  |  |  |  |
|  | L ITG | -48 | -54 | -12 | 6.60 |  |  |  |  |  |  |  |
|  | L Occipital | -24 | -63 | 39 | 6.42 |  |  |  |  |  |  |  |
|  | L Occipital | -24 | -90 | 18 | 5.33 |  |  |  |  |  |  |  |
|  | R Cerebelum | 9 | -75 | -27 | 5.25 |  |  |  |  |  |  |  |
|  | R AG | 33 | -57 | 45 | 5.20 |  |  |  |  |  |  |  |
|  | L Linual Gyrus | -9 | -75 | 9 | 4.98 |  |  |  |  |  |  |  |
|  | R Occipital | 30 | -72 | 30 | 4.47 |  |  |  |  |  |  |  |
|  | R Fusiform Gyrus | 39 | -60 | -15 | 3.63 |  |  |  |  |  |  |  |
|  | L IPL | -42 | -30 | 45 | 2.92 |  |  |  |  |  |  |  |
|  | L MPFC | -6 | 15 | 48 | 7.61 |  |  |  |  |  |  |  |
|  | R MFG | 45 | 27 | 27 | 6.71 |  |  |  |  |  |  |  |
|  | L Supramarginal Gyrus | -57 | -51 | 33 | -9.94 |  |  |  |  |  |  |  |
|  | R STG | 63 | -27 | 12 | -9.77 |  |  |  |  |  |  |  |
|  | L MPFC | -9 | 51 | 9 | -9.62 |  |  |  |  |  |  |  |
|  | L STG | -42 | -18 | 0 | -9.57 |  |  |  |  |  |  |  |
|  | R STG | 57 | -42 | 27 | -9.51 |  |  |  |  |  |  |  |
|  | L Insula | -39 | 6 | -6 | -9.33 |  |  |  |  |  |  |  |
|  | R MCC | 3 | -18 | 39 | -9.25 |  |  |  |  |  |  |  |
|  | R SupraMarginal Gyrus | 63 | -30 | 45 | -8.85 |  |  |  |  |  |  |  |
|  | L STG | -63 | -36 | 18 | -8.82 |  |  |  |  |  |  |  |
|  | R ITG | 48 | 3 | -39 | -8.47 |  |  |  |  |  |  |  |
|  | L MCC | -9 | -33 | 45 | -8.34 |  |  |  |  |  |  |  |
|  | R Cerebelum | 45 | -51 | -45 | -5.80 |  |  |  |  |  |  |  |
|  | L Cerebelum | -27 | -78 | -36 | -5.06 |  |  |  |  |  |  |  |
| Seed left IFG  Knowledge  vs.  Non-semantic | L Linual Gyrus | -21 | -93 | -9 | 8.01 |  | Seed left IFG  Knowledge  vs.  Non-semantic  Older > Young | R Pallidum | 12 | 3 | -3 | -4.65 |
|  | R Fusiform Gyrus | 24 | -87 | -9 | 7.70 |  |  | R IFG (pars opercularis) | 42 | 6 | 12 | -4.33 |
|  | L ITG | -45 | -48 | -15 | 3.60 |  |  | R Rolandic Operculum | 45 | -18 | 21 | -3.27 |
|  | L MPFC | -6 | 15 | 51 | 7.25 |  |  | R Insula | 36 | 15 | -9 | -3.23 |
|  | L IFG (pars triangularis) | -45 | 18 | 24 | 7.11 |  |  |  |  |  |  |  |
|  | L Precentral Gyrus | -51 | 0 | 48 | 3.84 |  |  |  |  |  |  |  |
|  | L MCC | -9 | -36 | 45 | -14.65 |  |  |  |  |  |  |  |
|  | L SFG | -27 | 33 | 39 | -13.55 |  |  |  |  |  |  |  |
|  | R MPFC | 9 | 51 | 3 | -13.54 |  |  |  |  |  |  |  |
|  | L Occipital | -42 | -84 | 30 | -13.52 |  |  |  |  |  |  |  |
|  | L Cuneus | -12 | -66 | 27 | -13.35 |  |  |  |  |  |  |  |
|  | R MCC | 9 | -27 | 42 | -13.22 |  |  |  |  |  |  |  |
|  | R SupraMarginal Gyrus | 60 | -45 | 36 | -13.08 |  |  |  |  |  |  |  |
|  | L SupraMarginal Gyrus | -57 | -45 | 36 | -12.66 |  |  |  |  |  |  |  |
|  | R ITG | 57 | -60 | 0 | -12.33 |  |  |  |  |  |  |  |
|  | R MTG | 45 | -72 | 24 | -12.17 |  |  |  |  |  |  |  |
|  | L MTG | -45 | -66 | 12 | -11.74 |  |  |  |  |  |  |  |
|  | L SFG | -21 | 15 | 57 | -11.70 |  |  |  |  |  |  |  |
|  | R MFG | 30 | 27 | 42 | -11.67 |  |  |  |  |  |  |  |
|  | L Fusiform Gyrus | -27 | -48 | -9 | -11.39 |  |  |  |  |  |  |  |
| Seed right IFG  Control  vs.  Non-semantic | L IFG (pars triangularis) | -45 | 21 | 24 | 8.81 |  | Seed right IFG  Control  vs.  Non-semantic  Older > Young | NA |  |  |  |  |
|  | L MPFC | -3 | 15 | 48 | 8.58 |  |  |  |  |  |  |  |
|  | L Insula | -30 | 21 | 3 | 7.20 |  |  |  |  |  |  |  |
|  | L MFG | -30 | 3 | 57 | 5.89 |  |  |  |  |  |  |  |
|  | L IFG (pars orbitalis) | -42 | 39 | 0 | 4.41 |  |  |  |  |  |  |  |
|  | L Linual Gyrus | -18 | -93 | -9 | 8.54 |  |  |  |  |  |  |  |
|  | R Occipital | 27 | -93 | -6 | 8.23 |  |  |  |  |  |  |  |
|  | L Occipital | -30 | -69 | 36 | 7.83 |  |  |  |  |  |  |  |
|  | L ITG | -39 | -42 | -21 | 7.44 |  |  |  |  |  |  |  |
|  | L SPL | -24 | -75 | 57 | 6.36 |  |  |  |  |  |  |  |
|  | L Thalamus | -9 | -12 | 6 | 4.55 |  |  |  |  |  |  |  |
|  | R Caudate | 9 | 12 | 9 | 4.41 |  |  |  |  |  |  |  |
|  | L Caudate | -12 | 9 | 6 | 4.07 |  |  |  |  |  |  |  |
|  | R IFG (pars triangularis) | 45 | 15 | 30 | 6.28 |  |  |  |  |  |  |  |
|  | R MFG | 30 | 0 | 51 | 4.61 |  |  |  |  |  |  |  |
|  | R Postcentral Gyrus | 45 | -27 | 45 | 3.78 |  |  |  |  |  |  |  |
|  | R Supramarginal Gyrus | 57 | -36 | 33 | -8.46 |  |  |  |  |  |  |  |
|  | R STG | 63 | -39 | 12 | -8.27 |  |  |  |  |  |  |  |
|  | R Cerebelum | 12 | -42 | -12 | -5.90 |  |  |  |  |  |  |  |
|  | L STG | -42 | -6 | -9 | -7.60 |  |  |  |  |  |  |  |
|  | L Supramarginal Gyrus | -60 | -39 | 33 | -6.46 |  |  |  |  |  |  |  |
|  | L STG | -45 | -36 | 12 | -5.75 |  |  |  |  |  |  |  |
|  | R MCC | 12 | -33 | 45 | -5.76 |  |  |  |  |  |  |  |
|  | R Postcentral Gyrus | 15 | -45 | 66 | -5.22 |  |  |  |  |  |  |  |
|  | L Postcentral Gyrus | -21 | -42 | 69 | -4.57 |  |  |  |  |  |  |  |
| Seed right IFG  Knowledge  vs.  Non-semantic | L MPFC | -6 | 12 | 51 | 9.82 |  | Seed right IFG  Knowledge  vs.  Non-semantic  Older > Young | R Olfactory cortex | 9 | 9 | -12 | -4.40 |
|  | L Linual Gyrus | -21 | -90 | -9 | 8.92 |  |  | R MPFC | 9 | 57 | 24 | -4.25 |
|  | L IFG (pars triangularis) | -42 | 15 | 24 | 8.33 |  |  | R MPFC | 6 | 36 | 0 | -4.07 |
|  | L IFG (pars orbitalis) | -30 | 24 | 0 | 7.01 |  |  | L MPFC | -12 | 48 | 9 | -4.02 |
|  | L Precentral Gyrus | -51 | 0 | 45 | 6.51 |  |  | R MPFC | 3 | 63 | -9 | -4.01 |
|  | L ITG | -45 | -45 | -18 | 5.98 |  |  | R Caudate | 15 | 15 | 9 | -3.09 |
|  | R Fusiform Gyrus | 24 | -87 | -9 | 9.06 |  |  | L PCC | 0 | -45 | 27 | -3.93 |
|  | R Fusiform Gyrus | 39 | -63 | -12 | 4.72 |  |  | L Calcarine Gyrus | -9 | -54 | 9 | -3.50 |
|  | R AG | 30 | -57 | 48 | 4.59 |  |  | R Insula | 39 | -12 | 12 | -3.84 |
|  | R Cerebelum | 12 | -72 | -24 | 4.54 |  |  | R STG | 42 | -33 | 21 | -3.65 |
|  | R Cerebelum | 36 | -42 | -24 | 3.80 |  |  | R Rolandic Operculum | 69 | -12 | 15 | -3.61 |
|  | R IFG (pars triangularis) | 42 | 12 | 30 | 5.38 |  |  |  |  |  |  |  |
|  | L PHG | -30 | -42 | -9 | -8.40 |  |  |  |  |  |  |  |
|  | R Supramarginal Gyrus | 60 | -39 | 36 | -7.52 |  |  |  |  |  |  |  |
|  | R ITG | 60 | -60 | 0 | -7.47 |  |  |  |  |  |  |  |
|  | L Calcarine Gyrus | -21 | -57 | 15 | -7.26 |  |  |  |  |  |  |  |
|  | L MCC | -9 | -39 | 45 | -7.23 |  |  |  |  |  |  |  |
|  | R Fusiform Gyrus | 30 | -42 | -9 | -7.07 |  |  |  |  |  |  |  |
|  | L Insula | -39 | -9 | -3 | -6.78 |  |  |  |  |  |  |  |
|  | R MTG | 42 | -69 | 27 | -6.65 |  |  |  |  |  |  |  |
|  | R Calcarine Gyrus | 24 | -54 | 15 | -6.46 |  |  |  |  |  |  |  |
|  | R MPFC | 9 | 42 | -3 | -6.28 |  |  |  |  |  |  |  |
|  | L Precuneus | -9 | -63 | 30 | -5.74 |  |  |  |  |  |  |  |
|  | R MTG | 60 | -18 | -15 | -5.25 |  |  |  |  |  |  |  |
|  | R ITG | 54 | -6 | -30 | -4.99 |  |  |  |  |  |  |  |

Note: AG, angular gyrus; ITG, inferior temporal gyrus; IFG, inferior frontal gyrus; IPL, Inferior Parietal Lobule; MCC, middle cingulate cortex; MTG, middle temporal gyrus; MFG, middle frontal gyrus; MPFC, medial prefrontal cortex; PCC posterior cingulate cortex; PHG, parahippocampal gyrus; SFG, superior frontal gyrus; SPL, superior parietal lobule; STG, superior temporal gyrus.

**Table S3. Peak MNI coordinates for the results of the whole-brain PPI analysis with left ATL and right ATL as seeds.**

|  | Effects across two age groups | | | | |  |  | Age group differences | | | | |
| --- | --- | --- | --- | --- | --- | --- | --- | --- | --- | --- | --- | --- |
|  | Region | x | y | z | t value |  |  | Region | x | y | z | t value |
| Seed left ATL  Control  vs.  Non-semantic | L MPFC | 0 | 18 | 48 | 8.18 |  | Seed left ATL  Control  vs.  Non-semantic  Older > Young | R STG | 63 | -45 | 18 | 5.44 |
|  | L IFG (pars triangularis) | -45 | 15 | 27 | 7.55 |  |  | R Medial Temporal Pole | 51 | 12 | -27 | 4.43 |
|  | L Insula | -27 | 21 | 3 | 6.85 |  |  | R Supramarginal Gyrus | 60 | -27 | 30 | 4.24 |
|  | L MFG | -27 | 3 | 54 | 5.67 |  |  | R STG | 60 | -21 | 6 | 3.77 |
|  | L Middle Orbital Gyrus | -48 | 42 | 3 | 5.36 |  |  | R ACC | 6 | 51 | 15 | 5.05 |
|  | R Fusiform Gyrus | 33 | -84 | -9 | 7.06 |  |  | L MPFC | -12 | 51 | 0 | 4.94 |
|  | L Fusiform Gyrus | -33 | -84 | -9 | 6.81 |  |  | R SFG | 18 | 36 | 36 | 3.66 |
|  | L ITG | -48 | -51 | -15 | 6.58 |  |  | L MPFC | -6 | 63 | 33 | 3.26 |
|  | R AG | 30 | -63 | 48 | 6.47 |  |  | L ACC | 0 | 30 | 15 | 3.24 |
|  | R MFG | 45 | 30 | 24 | 6.38 |  |  | L Insula | -42 | -12 | 0 | 4.98 |
|  | R MFG | 33 | 0 | 54 | 4.88 |  |  | L Supramarginal Gyrus | -57 | -42 | 36 | 4.46 |
|  | R MFG | 33 | 51 | 12 | 4.33 |  |  | L Hippocampus | -24 | -12 | -12 | 4.11 |
|  | R Supramarginal Gyrus | 63 | -33 | 36 | -7.07 |  |  | L MTG | -51 | -3 | -21 | 4.00 |
|  | R STG | 63 | -42 | 15 | -4.84 |  |  | L Precuneus | -9 | -63 | 30 | 3.98 |
|  | R MTG | 57 | -60 | 3 | -3.88 |  |  | R Precuneus | 15 | -54 | 24 | 3.58 |
|  | L Precuneus | -15 | -57 | 27 | -6.26 |  |  | R MCC | 6 | -24 | 42 | 3.55 |
|  | R MCC | 9 | -27 | 42 | -5.28 |  |  |  |  |  |  |  |
|  | R Postcentral Gyrus | 15 | -45 | 66 | -4.11 |  |  |  |  |  |  |  |
|  | L Precuneus | -12 | -45 | 69 | -3.67 |  |  |  |  |  |  |  |
|  | R Temporal Pole | 39 | 3 | -15 | -6.67 |  |  |  |  |  |  |  |
|  | R ITG | 51 | 3 | -39 | -4.26 |  |  |  |  |  |  |  |
|  | R Insula | 36 | 6 | 12 | -4.15 |  |  |  |  |  |  |  |
|  | R MTG | 57 | -21 | -12 | -3.79 |  |  |  |  |  |  |  |
|  | R STG | 57 | -3 | 3 | -3.46 |  |  |  |  |  |  |  |
|  | L Insula | -36 | 6 | 12 | -5.80 |  |  |  |  |  |  |  |
|  | L STG | -54 | -6 | 3 | -4.17 |  |  |  |  |  |  |  |
|  | L STG | -39 | -42 | 21 | -3.95 |  |  |  |  |  |  |  |
|  | L MTG | -48 | -60 | 21 | -3.48 |  |  |  |  |  |  |  |
|  | L SFG | -18 | 36 | 36 | -5.37 |  |  |  |  |  |  |  |
|  | R ACC | 6 | 33 | 9 | -5.05 |  |  |  |  |  |  |  |
|  | L MPFC | -9 | 51 | 0 | -4.89 |  |  |  |  |  |  |  |
|  | R SFG | 21 | 39 | 36 | -3.98 |  |  |  |  |  |  |  |
| Seed left ATL  Knowledge  vs.  Non-semantic | L Fusiform Gyrus | -33 | -78 | -12 | 8.41 |  | Seed left ATL  Knowledge  vs.  Non-semantic  Older > Young | L MPFC | -12 | 51 | 3 | 4.56 |
|  | R Occipital | 36 | -81 | -9 | 7.37 |  |  | R MPFC | 15 | 45 | 0 | 3.96 |
|  | L ITG | -48 | -51 | -15 | 6.85 |  |  | R MPFC | 6 | 54 | 21 | 3.80 |
|  | L Occipital | -27 | -87 | 9 | 6.80 |  |  | L Precuneus | -9 | -60 | 27 | 3.85 |
|  | L Occipital | -24 | -69 | 36 | 6.24 |  |  | R Cingulum | 15 | -45 | 36 | 3.49 |
|  | L MPFC | -6 | 18 | 48 | 7.23 |  |  |  |  |  |  |  |
|  | L IFG (pars triangularis) | -42 | 9 | 30 | 6.77 |  |  |  |  |  |  |  |
|  | L Insula | -30 | 21 | -3 | 6.48 |  |  |  |  |  |  |  |
|  | L IFG (pars orbitalis) | -45 | 39 | 0 | 6.42 |  |  |  |  |  |  |  |
|  | L Precentral Gyrus | -54 | 0 | 48 | 5.89 |  |  |  |  |  |  |  |
|  | L IFG (pars triangularis) | -48 | 15 | 6 | 5.37 |  |  |  |  |  |  |  |
|  | R IFG (pars triangularis) | 45 | 30 | 21 | 5.30 |  |  |  |  |  |  |  |
|  | L Precuneus | -9 | -63 | 30 | -6.71 |  |  |  |  |  |  |  |
|  | L PHG | -30 | -39 | -6 | -6.11 |  |  |  |  |  |  |  |
|  | L MCC | -9 | -33 | 45 | -5.73 |  |  |  |  |  |  |  |
|  | R Precuneus | 6 | -48 | 21 | -4.79 |  |  |  |  |  |  |  |
|  | R Supramarginal Gyrus | 63 | -36 | 36 | -5.77 |  |  |  |  |  |  |  |
|  | R AG | 42 | -66 | 27 | -4.28 |  |  |  |  |  |  |  |
|  | R MTG | 63 | -60 | 3 | -4.18 |  |  |  |  |  |  |  |
|  | R STG | 45 | -33 | 12 | -3.81 |  |  |  |  |  |  |  |
|  | L SFG | -27 | 33 | 39 | -5.66 |  |  |  |  |  |  |  |
|  | L MPFC | -9 | 51 | 0 | -5.64 |  |  |  |  |  |  |  |
|  | R Insula | 42 | -3 | -9 | -4.95 |  |  |  |  |  |  |  |
|  | R PHG | 33 | -36 | -9 | -4.90 |  |  |  |  |  |  |  |
|  | L Occipital | -45 | -81 | 30 | -5.14 |  |  |  |  |  |  |  |
|  | L MTG | -54 | -60 | 24 | -4.73 |  |  |  |  |  |  |  |
|  | L Supramarginal Gyrus | -60 | -33 | 30 | -4.07 |  |  |  |  |  |  |  |
| Seed right ATL  Control  vs.  Non-semantic | L IFG (pars opercularis) | -42 | 3 | 27 | 6.21 |  | Seed right ATL  Control  vs.  Non-semantic  Older > Young | R STG | 60 | -48 | 21 | 5.48 |
|  | L IFG (pars triangularis) | -45 | 30 | 15 | 4.96 |  |  | R MTG | 54 | -3 | -18 | 5.35 |
|  | L Precentral Gyrus | -60 | 6 | 36 | 4.96 |  |  | R Supramarginal Gyrus | 63 | -30 | 39 | 4.35 |
|  | L MFG | -24 | 0 | 54 | 4.93 |  |  | R MTG | 69 | -12 | -6 | 4.17 |
|  | R IFG (pars triangularis) | 45 | 33 | 18 | 6.11 |  |  | R MCC | 9 | -24 | 42 | 5.14 |
|  | R IFG (pars triangularis) | 42 | 9 | 27 | 4.64 |  |  | R Precuneus | 12 | -60 | 27 | 4.91 |
|  | L IPL | -36 | -39 | 39 | 5.77 |  |  | L Precuneus | -6 | -69 | 36 | 4.77 |
|  | L Occipital | -27 | -69 | 36 | 5.54 |  |  | L MCC | -12 | -48 | 42 | 4.18 |
|  | R Supramarginal Gyrus | 42 | -36 | 42 | 5.36 |  |  | R Fusiform Gyrus | 30 | -48 | -9 | 4.33 |
|  | L ITG | -45 | -60 | -9 | 5.31 |  |  | R PHG | 30 | -27 | -12 | 3.06 |
|  | L MPFC | 3 | 15 | 48 | 5.29 |  |  | L STG | -63 | -30 | 9 | 4.23 |
|  | L STG | -66 | -30 | 18 | -2.67 |  |  | L MTG | -60 | -54 | 18 | 4.08 |
|  | R PCC | 9 | -51 | 27 | -4.60 |  |  | R MPFC | 3 | 48 | -3 | 3.79 |
|  | R MCC | 6 | -24 | 39 | -3.15 |  |  |  |  |  |  |  |
|  | R MPFC | 9 | 57 | 3 | -4.50 |  |  |  |  |  |  |  |
|  | R SFG | 15 | 30 | 54 | -4.33 |  |  |  |  |  |  |  |
|  | R SFG | 15 | 57 | 39 | -3.71 |  |  |  |  |  |  |  |
|  | L MPFC | -9 | 51 | -6 | -3.33 |  |  |  |  |  |  |  |
|  | R Supramarginal Gyrus | 69 | -42 | 33 | -3.81 |  |  |  |  |  |  |  |
|  | R MTG | 57 | -54 | 12 | -3.81 |  |  |  |  |  |  |  |
|  | R Rolandic Operculum | 48 | -33 | 27 | -3.45 |  |  |  |  |  |  |  |
| Seed right ATL  Knowledge  vs.  Non-semantic | L IPL | -33 | -39 | 42 | 4.53 |  | Seed right ATL  Knowledge  vs.  Non-semantic  Older > Young | NA |  |  |  |  |
|  | L Fusiform Gyrus | -45 | -66 | -15 | 4.39 |  |  |  |  |  |  |  |
|  | L Occipital | -21 | -66 | 39 | 4.33 |  |  |  |  |  |  |  |
|  | L Occipital | -36 | -87 | 0 | 3.65 |  |  |  |  |  |  |  |
|  | L Cerebelum | -39 | -42 | -24 | 2.89 |  |  |  |  |  |  |  |
|  | R Occipital | 30 | -69 | 33 | 4.24 |  |  |  |  |  |  |  |
|  | R Occipital | 39 | -84 | -6 | 3.88 |  |  |  |  |  |  |  |
|  | R Calcarine Gyrus | 27 | -78 | 15 | 3.38 |  |  |  |  |  |  |  |
|  | R Fusiform Gyrus | 39 | -57 | -15 | 3.18 |  |  |  |  |  |  |  |
|  | L AG | -57 | -66 | 30 | -4.29 |  |  |  |  |  |  |  |
|  | L Occipital | -39 | -84 | 36 | -3.41 |  |  |  |  |  |  |  |

Note: ACC, anterior cingulate cortex; AG, angular gyrus; IFG, inferior frontal gyrus; IPL, inferior parietal lobule; ITG, inferior temporal gyrus; MCC, middle cingulate cortex; MFG, middle frontal gyrus; MPFC, medial prefrontal cortex; MTG, middle temporal gyrus; PCC posterior cingulate cortex; PHG, parahippocampal gyrus; SFG, superior frontal gyrus; STG, superior temporal gyrus.

**Table S4. Peak MNI coordinates for the results of the whole-brain PPI analysis for semantic task effects, with each of the ROIs as the seed.**

|  | Effects across two age groups | | | | |  |  | Age group differences | | | | |
| --- | --- | --- | --- | --- | --- | --- | --- | --- | --- | --- | --- | --- |
|  | Region | x | y | z | t value |  |  | Region | x | y | z | t value |
| Seed left IFG  Control  vs.  Knowledge | L Occipital | -36 | -78 | 27 | 14.55 |  | Seed left IFG  Control  vs.  Knowledge  Older > Young | R Temporal Pole | 33 | 18 | -39 | 4.62 |
|  | L Cerebelum | -12 | -75 | -9 | 14.10 |  |  | R MPFC | 12 | 51 | 18 | 3.83 |
|  | L MFG | -24 | 12 | 57 | 12.94 |  |  | R MPFC | 9 | 63 | -3 | 3.48 |
|  | R Precuneus | 3 | -60 | 45 | 12.66 |  |  | L SFG | -15 | 51 | 12 | 3.47 |
|  | R Occipital | 39 | -75 | 30 | 12.36 |  |  | L Rectal Gyrus | -3 | 42 | -15 | 2.93 |
|  | R MFG | 27 | 15 | 54 | 12.13 |  |  | L Linual Gyrus | -3 | -69 | 9 | -3.53 |
|  | L Calcarine Gyrus | 0 | -66 | 18 | 11.82 |  |  | R Linual Gyrus | 21 | -63 | 3 | -3.29 |
|  | L Fusiform Gyrus | -30 | -39 | -15 | 11.17 |  |  | L Cuneus | -12 | -84 | 21 | -3.25 |
|  | R Linual Gyrus | 9 | -72 | -3 | 11.16 |  |  |  |  |  |  |  |
|  | L IFG (pars triangularis) | -45 | 39 | 18 | 10.92 |  |  |  |  |  |  |  |
|  | L ITG | -54 | -60 | -9 | 10.77 |  |  |  |  |  |  |  |
|  | L Linual Gyrus | -15 | -48 | -3 | 10.53 |  |  |  |  |  |  |  |
|  | R Linual Gyrus | 12 | -48 | 3 | 10.17 |  |  |  |  |  |  |  |
|  | R ITG | 54 | -48 | -9 | 9.49 |  |  |  |  |  |  |  |
|  | L MFG | -39 | 27 | 33 | 9.39 |  |  |  |  |  |  |  |
|  | R Fusiform Gyrus | 30 | -39 | -15 | 8.63 |  |  |  |  |  |  |  |
|  | R IFG (pars triangularis) | 42 | 27 | 33 | 8.58 |  |  |  |  |  |  |  |
|  | R MFG | 42 | 42 | 15 | 8.47 |  |  |  |  |  |  |  |
|  | R Supramarginal Gyrus | 48 | -36 | 48 | 8.41 |  |  |  |  |  |  |  |
|  | L IPL | -36 | -48 | 45 | 8.30 |  |  |  |  |  |  |  |
|  | R Cuneus | 18 | -84 | 27 | 7.98 |  |  |  |  |  |  |  |
|  | L Precuneus | -18 | -66 | 39 | 7.97 |  |  |  |  |  |  |  |
|  | R Middle Orbital Gyrus | 27 | 36 | -12 | 7.93 |  |  |  |  |  |  |  |
|  | R SFG | 27 | 57 | 9 | 7.86 |  |  |  |  |  |  |  |
|  | L SFG | -24 | 60 | 12 | 7.80 |  |  |  |  |  |  |  |
| Seed right IFG  Control  vs.  Knowledge | L Occipital | -36 | -81 | 33 | 6.84 |  | Seed right IFG  Control  vs.  Knowledge  Older > Young | NA |  |  |  |  |
|  | L AG | -42 | -63 | 42 | 4.57 |  |  |  |  |  |  |  |
|  | L MTG | -51 | -54 | 21 | 4.36 |  |  |  |  |  |  |  |
|  | L IPL | -42 | -54 | 60 | 3.35 |  |  |  |  |  |  |  |
|  | L Precuneus | -3 | -60 | 39 | 6.41 |  |  |  |  |  |  |  |
|  | L Fusiform Gyrus | -27 | -42 | -12 | 5.86 |  |  |  |  |  |  |  |
|  | L Calcarine Gyrus | -9 | -57 | 12 | 5.86 |  |  |  |  |  |  |  |
|  | L Cerebelum | -9 | -72 | -6 | 5.69 |  |  |  |  |  |  |  |
|  | R Cerebelum | 12 | -72 | -9 | 5.50 |  |  |  |  |  |  |  |
|  | L Calcarine Gyrus | 3 | -81 | 15 | 4.18 |  |  |  |  |  |  |  |
|  | R Cerebelum | 15 | -84 | -27 | 3.03 |  |  |  |  |  |  |  |
|  | R MFG | 24 | 27 | 42 | 6.06 |  |  |  |  |  |  |  |
|  | R SFG | 24 | 57 | 12 | 4.67 |  |  |  |  |  |  |  |
|  | L MPFC | 3 | 42 | 30 | 4.46 |  |  |  |  |  |  |  |
|  | R MPFC | 6 | 60 | -3 | 4.25 |  |  |  |  |  |  |  |
|  | R MFG | 42 | 51 | 27 | 3.83 |  |  |  |  |  |  |  |
|  | L ACC | -9 | 42 | 12 | 3.59 |  |  |  |  |  |  |  |
|  | L MFG | -21 | 15 | 51 | 5.94 |  |  |  |  |  |  |  |
|  | L IFG (pars triangularis) | -42 | 39 | 18 | 3.41 |  |  |  |  |  |  |  |
|  | R AG | 48 | -63 | 36 | 5.86 |  |  |  |  |  |  |  |
|  | R IPL | 45 | -60 | 57 | 4.48 |  |  |  |  |  |  |  |
|  | R Postcentral Gyrus | 48 | -33 | 48 | 3.15 |  |  |  |  |  |  |  |
|  | L ITG | -60 | -57 | -6 | 5.63 |  |  |  |  |  |  |  |
|  | R ITG | 57 | -48 | -9 | 5.32 |  |  |  |  |  |  |  |
|  | R ITG | 66 | -15 | -24 | 4.88 |  |  |  |  |  |  |  |
| Seed left ATL  Control  vs.  Knowledge | R Precuneus | 3 | -63 | 45 | 4.19 |  | Seed left ATL  Control  vs.  Knowledge  Older > Young | NA |  |  |  |  |
|  | L Calcarine Gyrus | 0 | -66 | 18 | 3.95 |  |  |  |  |  |  |  |
| Seed right ATL  Control  vs.  Knowledge | NA |  |  |  |  |  | Seed right ATL  Control  vs.  Knowledge  Older > Young | R PCC | 6 | -24 | 24 | 4.44 |
|  |  |  |  |  |  |  |  | R Precuneus | 18 | -63 | 39 | 3.90 |

Note: ACC, anterior cingulate cortex; AG, angular gyrus; IFG, inferior frontal gyrus; IPL, inferior parietal lobule; ITG, inferior temporal gyrus; MFG, middle frontal gyrus; MTG, middle temporal gyrus; MPFC, medial prefrontal cortex; PCC, posterior cingulate cortex; SFG, superior frontal gyrus.
